# A single-nuclei RNA sequencing study of Mendelian and sporadic AD in the human brain

**DOI:** 10.1101/593756

**Authors:** Jorge L. Del-Aguila, Zeran Li, Umber Dube, Kathie A. Mihindukulasuriya, John P Budde, Maria Victoria Fernandez, Laura Ibanez, Joseph Bradley, Fengxian Wang, Kristy Bergmann, Richard Davenport, John C. Morris, David M. Holtzman, Richard J. Perrin, Bruno A. Benitez, Joseph Dougherty, Carlos Cruchaga, Oscar Harari

**Author notes:** To whom correspondence should be addressed: Carlos Cruchaga, PhD, Professor Dep. of Psychiatry, Genetics and Neurology, Washington University School of Medicine, Campus Box 8134 | 425 S. Euclid Ave |BJC Institute of Health |Office: 9607 | St. Louis, MO, 63110, |, Oscar Harari, PhD, Assistant Professor, Department of Psychiatry, Washington University School of Medicine, Campus Box 8134 | 425 S. Euclid Ave |BJC Institute of Health |Office: 9615 | St. Louis, MO, 63110, |.

## Abstract

Alzheimer Disease (AD) is the most common form of dementia. This neurodegenerative disorder is associated with neuronal death and gliosis heavily impacting the cerebral cortex. AD has a substantial but heterogeneous genetic component, presenting both Mendelian and complex genetic architectures. Using bulk RNA-seq from parietal lobes and deconvolution methods, we previously reported that brains exhibiting different AD genetic architecture exhibit different cellular proportions. Here, we sought to directly investigate AD brain changes in cell proportion and gene expression using single cell resolution. To do so, we generated unsorted single-nuclei RNA-sequencing data from brain tissue. We leveraged tissue donated from a carrier of a Mendelian genetic mutation and two family members who suffer from AD, but do not have the same mutation. We evaluated alternative alignment approaches to maximize the titer of reads, genes and cells with high quality. In addition, we employed distinct clustering strategies to determine the best approach to identify cell clusters that reveal neuronal and glial cell types and avoid artifacts such as sample and batch effects. We propose an approach to cluster cells that reduces biases and enable further analyses. We identified distinct types of neurons, both excitatory and inhibitory, and glial cells, including astrocytes, oligodendrocytes, and microglia among others. In particular, we identified a reduced proportion of excitatory neurons in the Mendelian mutation carrier, but a similar distribution of inhibitory neurons. Furthermore, we investigated whether single-nuclei RNA-seq from human brains recapitulate the expression profile of Disease Associated Microglia (DAM) discovered in mouse models. We also determined that when analyzing human single-nuclei data it is critical to control for biases introduced by donor specific expression profiles. In conclusion, we propose a collection of best practices to generate a highly-detailed molecular cell atlas of highly informative frozen tissue stored in brain banks. Importantly, we have developed a new web application to make this unique single-nuclei molecular atlas publicly available.

## Introduction

Alzheimer Disease (AD) is a neurodegenerative disorder characterized by the presence of amyloid Aβ plaques and neurofibrillary tangles (hyperphosphorylated tau deposits) in the brain [19]. AD is also associated with neuronal death and gliosis specifically in the cerebral cortex. AD has a substantial but heterogeneous genetic component. While carriers of mutations in the *amyloid-beta precursor protein* (*APP*) and *Presenilin* genes (*PSEN1 and PSEN2*) [8, 30] show Mendelian inheritance patterns, the majority of the AD cases (90-95%) present a complex genetic architecture (Sporadic AD), with many genetic factor contributing to risk. Recently, we studied the cellular population structure of AD brains of carriers of Mendelian mutations in *APP, PSEN1* and *PSEN2*, sporadic ADs, and compared them to neuropathology free controls [22]. To do so, we generated bulk RNA-seq from the parietal lobe and analyzed it using an optimized digital deconvolution method [22] to infer broad proportions of neurons, astrocytes, oligodendrocytes and microglia. Importantly, we identified that brains of carriers of Mendelian mutations have a specific distribution of neurons that differs from that of sporadic AD [22]. However, bulk RNA-seq and deconvolution approaches do not provide a detailed context of expression profiles at the cellular level, which hampers the identification of which neuronal subtypes [20] are the most vulnerable to the AD pathogenesis.

Single-cell RNA-seq (scRNA-seq) provides the opportunity to generate the detailed transcriptomic cell profiling required to address the drawbacks of bulk RNA-seq [1, 27, 37]. However, performing scRNA-seq requires cell re-suspension and library preparation needs to be done from fresh tissue. This requirement prohibits the study of highly informative frozen human brain tissue stored in brain banks. Single-nuclei RNAseq (snuclRNA-seq) is an alternative to this methodology. Studies of mouse neural progenitor cells shows that only small differences are present between total cellular and nuclear RNA profiles [11]. We sought to investigate the feasibility of generating and employing unsorted snuclRNA-seq to analyze banked brains from related individuals, which exhibit Mendelian or sporadic genetic architectures of AD. We hypothesize that cell-type expression profiling data from these brains will provide a unique resource to: i) obtain a highly detailed map of cell composition in human brains of AD; ii) determine cellular population structure changes in common and specific to Mendelian and sporadic AD; and iii) characterize transcriptomic profile alterations in AD within each defined cell type.

We discovered that analyzing snuclRNA-seq from human brains has its own challenges, as most of the current tools and pipelines are set up to analyze scRNA-seq obtained from fresh tissue. When applying these to snculRNA-seq, we encountered artifacts and biases that constrained downstream analyses. Thus, we propose an approach to extend scRNA-seq analyzing methodology to snuclRNA-seq to identify clusters of nuclei that show trustworthy expression profiles, and which resemble distinct subtypes of neurons and glial cells, including oligodendrocytes, microglia, and astrocytes among others. In addition, these clusters have an even (fair) representation of nuclei from all of the donors, a property required by many single-nuclei downstream analyses. We then employed this approach to process the snuclRNA-seq from human frozen brains, and analyze the distribution of neuronal subtypes in brains with Mendelian and complex genetic architectures of AD.

Recently, a novel type of microglia has been proposed to be associated with neurodegeneration [18]. The Disease Associated Microglia (DAM) was identified generating scRNA-seq from sorted microglial cells from mouse models [18], and then validated using immunohistochemical staining of human brain tissue to demonstrate the colocalization of Aβ particles with a protein marker of DAM [18]. Here, we analyze whether the expression profile of human microglia detected by snuclRNA-seq recapitulate the DAM expression signature detected in mouse models.

Finally, our findings indicate that snuclRNA-seq provides a valuable resource that can improve our understanding of the sequence of regulatory events that control cell fate leading to AD. Therefore, we are making available the nuclei specific expression profile of the brains we analyzed through and interactive web-based application (http://ngi.pub/snuclRNA-seq) to provide full access to the research community to this resource. This is the first single-nuclei molecular atlas of AD brains carrying pathological mutations in *PSEN1* and related sporadic AD. We hope that this high-quality data will help elucidate and validate novel biological insights into AD and contribute to early diagnosis and therapeutic intervention.

## Methods

### 1. Samples

The Neuropathology Core of the Knight-Alzheimer’s Disease Research Center (Knight-ADRC) provided the parietal lobe tissue from postmortem brains for each sample. These samples were obtained with informed consent for research use and were approved by the institutional review board of Washington University in St. Louis. AD neuropathological changes were assessed according to the criteria of the National Institute on Aging-Alzheimer’s Association (NIA-AA). One AD patient is a carrier of the *PSEN1* p.A79V pathogenic mutation, while the two relatives do not carry any autosomal mutations (sporadic AD). The three donors are females with European-American female ancestry. Additional information, including clinical dementia rating (CDR), *APOE* genotype, postmortem interval (PMI), age at onset, and age at death are detailed in Table 1.

**Table 1:** Demographic Characteristics of samples.

|  | **Sample1** | **Sample2** | **Sample3** |
| --- | --- | --- | --- |
| **PSEN1 mutation** | Non-carrier | Non-carrier | Ala79Val |
| **Gender** | Female | Female | Female |
| **Age on set** | 87 | 81 | 76 |
| **Age at death** | 89 | 82 | 89 |
| **CDR** | 1 | 3 | 3 |
| **APOE** | □3/□4 | □3/□3 | □3/□4 |
| **PMI** | 9 | 17.2 | 8 |

### 2. Nuclei extraction and Library preparation

From the fresh frozen human parietal lobes, approximately 500mg of tissue were cut and weighed on dry ice using sterile disposable scalpels. The parietal tissue was homogenized in ice-cold homogenization buffer (0.25M Sucrose, 150 mM KCl, 5 mM MgCl2, 20 mM Tricine-KOH pH 7.8, 0.15mM spermine, 0.5mM spermidine, EDTA free protease inhibitor, and recombinant RNase inhibitors) with a glass-on-glass dounce homogenizer, 10 strokes with the A pestle, followed by 10 strokes of the B pestel. Homogenates were centrifuged for 5 min at 500 × g, at 4°C, to pellet the nuclear fraction. The nuclear fraction was mixed with an equal volume of 50% iodixanol and added on top of a 35% iodixanol solution for 30 min at 10,000 × g, at 4°C. After removal of the myelin layer from the top of the gradient, the nuclei were collected from the 30%-35% iodixanol interface. Nuclei were resuspended in nuclei wash and resuspension buffer (1.0% bovine serum albumin and recombinant RNase inhibitors in phosphate-buffered saline) and pelleted for 5 min at 500 × Gs, 4°C. Nuclei were passed through a 40 µM cell strainer to remove cell debris and large clumps. Nuclei concentration was manually determined using DAPI counterstaining and hemocytometer. Nuclei concentration was adjusted to 1,200 nuclei/µL and followed immediately by the 10x Genomics® Single Cell Protocol. We generated snuclRNA-seq libraries using the 10X Chromium V(D)J 5` chemistry for 10,000 cells per sample and sequenced 50,000 reads per cell from the three frozen human parietal lobes.

### 3. Sequencing Alignment and Data for Secondary Analysis

The CellRanger (v2.1.1 10XGenomics) software was employed to align the sequences and quantify gene expression. By default, this software quantifies expression for mature messenger RNA (mRNA) by counting reads aligned to exons as annotated in the human genome build GRCh38. However, the snuclRNA-seq profiles nuclear precursor mRNA (pre-mRNA), which include transcripts that have not completed splicing to remove introns. To capture all the information in the pre-mRNA, we aligned the reads to a custom “pre-mRNA” reference, that was generated as described by 10X genomics technical manual [1] (See Online Resources). In this way, the intronic reads from pre-mRNA are included in the final gene expression counts. We aligned and quantified gene expression using both mature and pre-mRNA references, and these reads were further cleaned QCed and compared.

### 4. Single-nuclei RNA-seq Cleaning

The gene expression matrices from all samples were combined in R independently for further processing using the Seurat (version 2.20, 2.30) pipeline. We processed the gene expression quantified for the pre-mRNA and mRNA references in parallel. We removed most of mitochondrial genes (<0.1%) and then kept genes expressed in three or more cells. We discarded cells with less than 1800 or more than 8000 genes expressed. To exclude multiplets, when a single droplet includes two or more nuclei, the top 0.5% of the distribution of nUMI (total number of Unique Molecular Identifiers detected in each cell) were removed. We also studied whether the number of genes expressed per nuclei followed a multi-modal distributions, under the assumption that glial cells could expressed a lower number of genes than neurons [29] This analysis was performed by the R package mixtools [3]. We normalized the data employing the *LogNormalize* function, which normalizes the gene expression measurements for each cell by the total expression; scales by a factor equal to the median counts of all genes and log-transforms the expression. Data regression was performed using the *ScaleData* function with nUMI, percent mitochondrial reads and sample origin as confounding factors.

### 5. Selection of Genes for cell clustering

The snuclRNA-seq expression data is highly dimensional, as thousands of nuclei are ascertained transcriptome-wide for each brain. However, as single cell methods are lossy, the quantification of snuclRNA-seq produces sparse data matrices, and the expression of each gene is not detected for each nuclei. The structure of the data complicates the clustering of nuclei into distinct cell types, and conventional similarity distances, such as Euclidean distance, have been claimed to be less reliable as the dimensions of the feature space increase [36]. However, only hundreds of genes are required to discriminate cell types. The selection of a representative set of genes to cluster nuclei is a key step in the processing of data, in which only the informative genes are filtered, in such a way that clustering of nuclei is computationally feasible. Given the importance of this processing step we evaluated alternative approaches, described below

#### 5.1 Classic Gene Set (CGS) from Pooling Subjects

The CGS method is the approach most commonly employed to select the most variable genes in scRNA-seq studies [7, 13]. The Seurat *FindVariableGenes* function performs this selection. We executed this method by using the default values and cut-offs (x.low.cutoff = 0.0125, x.high.cutoff = 3, y.cutoff = 0.8) and selected the most variable genes from the single nuclei expression from the three brains. We obtained 2,360 genes that were used to calculate 100 principal components (PCs). Then, we identified the optimal number of PCs for downstream analysis using heuristics, PC elbow plot, and JackStraw statistical tests. The *JackStraw* function randomly permutes a subset of data, and calculates projected PC scores for these genes. This analysis indicated that the first 65 PCs were sufficient for clustering the nuclei.

#### 5.2 Hicat Gene Markers

This method proposed by Taasic et al [32], employs known genes markers of cell types to generate an initial partition of the cells into a broad cell clusters. Later on, this initial partition can be subclustered to identify cell subtypes. For our analysis, we employed all nuclei that passed QC, using 118 known genes markers, that we collected from the literature (Online Resource Table S1) [2, 20, 24, 28, 29]. We calculated the first 100 PCs from the expression of these marker genes, as quantified using the pre-mRNA annotation (Seurat software). We selected the number of PCs using the same methodology as described in section 5.1 and selected the first 55 PCs for downstream analysis.

#### 5.3 Consensus Gene Set (ConGen)

The approach we propose, the Consensus Gene Set (ConGen), controls for biases and obtains clusters with even representation of all samples. We applied the Seurat *FindVariableGenes* with default selections and cut-offs values (x.low.cutoff = 0.0125, x.high.cutoff = 3, y.cutoff = 0.8), for each library sample and identified a set of highly variable genes for each brain sample whose expression was quantified using pre-mRNA annotation. The number of highly variable genes is 2,447; 2,354 and 1,972 for Sample1, Sample2 and Sample3, respectively. Then we identified the common set of genes that were highly variable among the samples using the R function intersection (N=1,434). These common genes were used to calculate 100 PCs from all of the samples (Seurat package). We selected the number of PCs using the same methodology described in section 5.1, and employed the first 25 PCs for downstream analysis.

### 6. Cell/Nuclei Clustering

The goal of this process is to group nuclei based on the similarities between their expression profiles. We used the Seurat function *FindClusters* to identify the clusters with a resolution parameter 0.6, and employed the *TSNEPlot* function to generate a visual representation of the clusters using T-distributed Stochastic Neighbor Embedding (tSNE). In addition, we corrected for drop-out events that lead to an exceedingly sparse depiction of the single cell transcriptome. Instead of removing genes containing missing values, which restricts the analysis to only highly expressed genes, we imputed missing gene expression values using *scImpute* [21]. After imputing gene expression, we clustered the nuclei again. Finally, we learned similarity relationships among the expression profile of the cells included in the clusters by using the function *BuildClusterTree*.

### 7. Coincidence Analysis

To compare the results of distinct clustering approaches and identify whether three different approaches were producing (dis)similar clusters, we implemented a coincidence analyses. We performed an exhaustive comparison of each cluster identified by one processing and clustering approach to all of the clusters identified by a second approach. In this way, we can determine how the cells are reorganized among different clusters, or in contrast to identify if cells are grouped coincidently together by distinct approaches. To do so, we calculate the normalized pointwise mutual information (*pmi*):

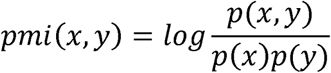

which quantifies the discrepancy between the probability of joint distribution of two clusters and their individual distributions. In addition, we calculated the Jaccard index, or similarity coefficient, which is defined as:

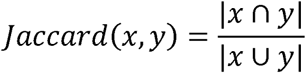

The Jaccard index evaluates the extent of the intersection of the nuclei in common between two clusters, corrected by the extent of the union of the nuclei assigned to the two clusters. To provide a global overview of the clusters’ coincidence, we plotted the pointwise comparisons and we represented the *pmi* by color and the *Jaccard index* by size of the point (Online Resource).

### 8. Entropy as a measure to evaluate donor evenness and biases in clustered nuclei

Having an even representation of nuclei from distinct samples in a cluster is critical to perform unbiased comparisons. Otherwise, clusters of nuclei with overrepresentation or underrepresentation of samples precludes many downstream analyses. We employed Shannon’s information theory entropy [31] as a quantitative measure to evaluate how even or biased the distribution of samples among the nuclei of any given cluster is. Specifically, entropy of a cluster is calculated as:

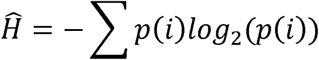

where *p(i)* is the probability of a nuclei belonging to subject *i* (Online Resources). The entropy tends to 0 when all of the cells from a cluster belong to a single subject, and is maximized when the distribution of subjects is perfectly uniform which is 1.58 for three samples (Online Resources).

### 9. Cluster annotation

We determined the brain cell types in each of the cluster by evaluating the expression of maker genes for neurons, astrocytes, oligodendrocytes, microglia, oligodendrocyte precursor cells and endothelial cells, usually employed in the literature [2, 20, 24, 28, 29] (Online Resource Table S2 and Table S3). We used the *DotPlot* function from the Seurat package to visualize the average expression of genes related to specific cell types. To determine the homogeny of brain samples analyzed, we also evaluated the expression of marker genes tagging distinct pyramidal layers for the excitatory neurons. We also looked for excitatory and inhibitory neurons [20].

### 10. Pseudo-temporal trajectories of microglia expression profiles

The gene expression for the microglia was analyzed using the R package TSCAN (version 1.7.0) to infer a pseudo-temporal path [16]. This method orders cells based on the transition of their transcriptomes, by initially clustering cells, then identifying similarity relationships among the clusters, inferring a Minimal Spanning Tree (MST), projecting cells onto tree edges and finally using generalized additive models to ascertain the functional relationship between the pseudo-time and gene expression. The raw expression data from microglia cells was loaded and pre-processed (TSCAN function *preprocess*) using default parameters. We then executed the *exprmclust* function with default parameters to learn the optimal number of clusters for microglia. Then, we employed the *tscan* function order to identify pseudo-time trajectories of the cells. Finally, we used the processed expression data and the pseudo-order to identify the genes associated with this ordering, using the *difftest* function.

### 11. Single-nuclei gene expression browser

We developed a web application to provide public access to the single-nuclei transcriptomic atlas. All of the snuclRNA-seq data that we generated and processed can be queried and visualized using a custom browser that we have developed using the R Shiny framework. This browser provides an input panel to query individual genes and additional parameters to customize the visualization. For each gene, the browser provides graphical information organized in three panels: i) The tSNSE projection of the nuclei is represented and colored by the clusters. We identified and annotated these clusters using Seurat *TSNEPlot* function. ii) A gene expression level for each nuclei is graphically represented using the same tSNE representation. Nuclei are colored according to expression level of the queried gene. We represent nuclei highly expressing the target gene in purple and nuclei with lower expression in grey (Seurat *FeaturePlot* function). iii) We show the cell-type specific differential expression (log fold change, p-value and adjusted p-value) of queried genes for Neurons, Astrocytes, Microglia, Oligodendrocyte, Oligodendrocyte Precursor Cells (OPC) and Endothelial Cells that was pre-computed beforehand. We first grouped clusters by cell type (Seurat *BuildClusterTree* function). As a result, we obtained a unique cluster that includes all of the neurons, while conserving clusters for each of the remaining cell types. Thus, the nuclei are partitioned into a total of six clusters. Then we calculated the differential expression for each gene among these six clusters using the Seurat *FindMarkers* function. The statistical significance was calculated using the non-parameteric Wilcoxon rank sum test to determined those genes that had a log foldchange >0.01. In this way we generated a broader comparison of the transcriptome. The adjusted p-value is adjusted using a Bonferroni multiple test correction. All of the source code for the snuclRNA-seq explorer and the precomputation of the differential expression is available in the GitHub repository (https://github.com/NeuroGenomicsAndInformatics/snuclRNA-seq).

### 12. Bulk RNA-seq processing

In parallel, we generated bulk RNA-seq data for the three brains that we analyzed, and followed the same QC and processing as we reported previously [22]. These data were mapped to mRNA reference for the human genome build GRCh38.

### 13. Ascertaining snuclRNA-seq quality

To control for the quality of the processed transcriptomic snuclRNA-seq, we compared the gene expression values from both alignments for each subject, and as a group, to their gene expression values from their bulk RNA-seq.

## Results

### Study Design

We collected snuclRNA-seq data from three European-American female donors. We employed two different references for alignment, pre-mRNA and mRNA, to quantify the gene expression for each sample donor. Each sample was accompanied by processing of a matching bulk RNA-seq sample using mRNA as reference. These steps allowed us to estimate how well snuclRNA-seq data can recapitulate the RNA-seq results from bulk samples. The gene expression matrices from all samples were combined for further processing. Three different approaches were tested to select the variable genes that were used to determine the principal components for the nuclei clustering. We processed each approach in parallel. We tested the evenness of each cluster for each approach as well as critical parameters such as resolution and coincident analysis. The approach with the best results was then imputed and used for further analyses including the pseudo-time trajectories of the cells and Disease Associated Microglia association. Finally, we prepared a browser using the R Shiny framework to visualize the snuclRNA-seq data that we produced.

### Clinical and Demographic Characteristics

The complexity and uniqueness of the cells types in the different regions and layers in the human brain was previously described [10, 15, 17, 20, 26, 33, 34], however the sample quality is diminished in postmortem samples. To obtain highly detailed maps of cell composition in AD brains with distinct genetic architecture and to characterize their transcriptomic profiles at a cellular level, we analyzed snuclRNA-seq from the parietal lobe [4, 6, 14, 23], for three subjects (Online Resource Figure S1). All three donors were European-American females, with an age of death ranging from 82 to 89 years and were members of the same family. One of the donors, Sample3, was a carrier of the *PSEN1* p.A79V (ADAD) while the other two donors, Sample1 and Sample2, presented a complex genetic architecture of AD. Neuropathology shown definitive AD and the time between death. Sample collection or post mortem interval (PMI) ranged between 8 to 17.2 hours (Table 1).

### Generation and quality evaluation of human brain tissue single-nuclei RNA-seq

We generated snuclRNA-seq libraries using the 10X Chromium for 10,000 cells per sample and sequenced 50,000 reads per cell from three frozen human parietal lobes. We employed two different alignment libraries for the annotation of the genome, pre-mRNA and mRNA, and quantified gene expression using both (Methods). We observed that when we aligned the reads to pre-mRNA, the number of reads, genes and cells increase significantly. For example, when using pre-mRNA as a reference, we observed a 36% increase in the number of nuclei (26,331 vs 19,302; χ^2^ test, p <= 2.2×10^−16^, Table 2), without affecting the number of identified transcripts (28,428 vs 25,465 counts). We also observed a 118.8% increase in the number of the median UMI counts per nuclei (t-test, p=1.5×10^−2^) and a 45.1% for the median genes per nuclei (t-test, p=5.0×10^−3^).

**Table 2:** Summary Statistics After Quality Control.

| **Alignment to Precursor-RNA (pre-RNA)** |  |  |  |  |
| --- | --- | --- | --- | --- |
|  | **Sample1** | **Sample2** | **Sample3** | **ALL** |
| Number of nuclei | 5,663 | 7,147 | 13,521 | 26,331 |
| Median UMI Counts per nuclei | 11,560 | 14,444 | 6,543 | 9,262 |
| Median Genes per nuclei | 4,006 | 5,064 | 2,852 | 3,642 |
| Total Genes Detected | 28,428 | 28,428 | 28,428 | 28,428 |
| **Alignment to mature-RNA** |  |  |  |  |
|  | **Sample1** | **Sample2** | **Sample3** | **ALL** |
| Number of nuclei | 4,493 | 7,197 | 7,612 | 19,302 |
| Median UMI Counts per nuclei | 3,960 | 7,162 | 2,464 | 4,234 |
| Median Genes per nuclei | 2,355 | 3,851 | 1,579 | 2,510 |
| Total Genes Detected | 25,469 | 25,469 | 25,469 | 25,469 |

To further evaluate the quality of snuclRNA-seq data, we compared the gene expression values from the two alignments “pre-mRNA” and “mRNA” in snuclRNA-seq to their gene expression values from their parallel bulk RNA-seq aligned to the mRNA reference data (Table 3). We observed a Pearson correlation of r^2^=91% between snuclRNA-seq with mRNA reference and bulk RNAseq (Figure 1A). On the other hand, the Pearson correlation of snuclRNA-seq with pre-mRNA reference against bulk RNA-seq was r^2^=86% (Figure 1B). These results may suggest that aligning to mRNA is a better approach, but it is important to note that the alignment to pre-mRNA increased the number of nuclei, Median UMI Counts per nuclei and Median Genes per nuclei significantly. In addition, the correlation we identified for the pre-mRNA is in line with the values previously reported for single-cell RNA-seq [35].

**Table 3:** Correlation Analysis between bulk RNA-seq and snuclRNA-seq after QC.

|  | **Reference**<br>**Alignment** | **Bulk RNA-seq** |
| --- | --- | --- |
|  |  | **Mature-RNA** |
| **snRNA-seq** | **Pre-RNA** | 0.91 |
|  | **Mature-RNA** | 0.86 |

All values are Pearson coefficient

**Table 4:**
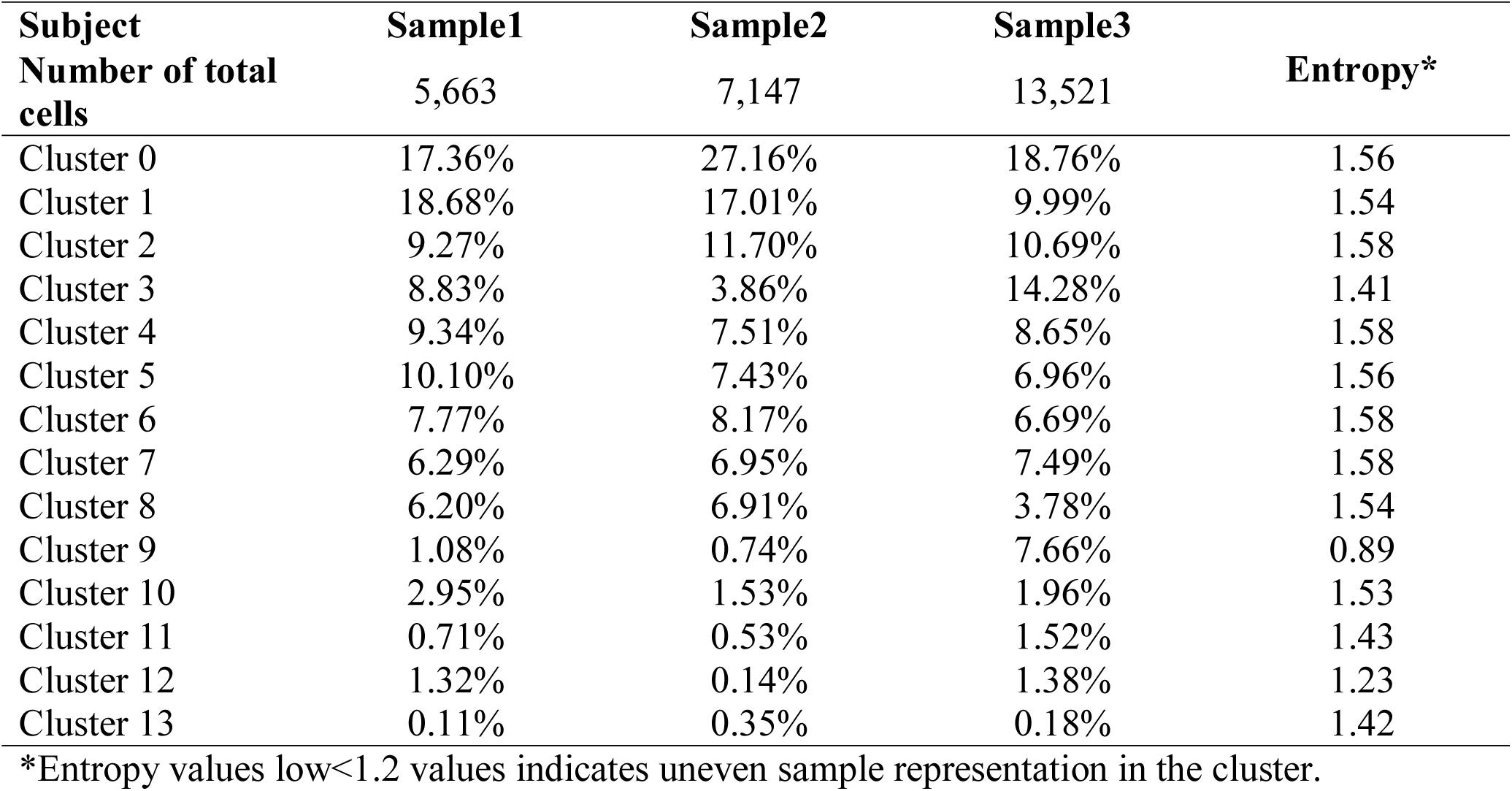
Number of cells for each subject in each cluster using imputed Consensus Gene Set data.

**Table 5:** Number of cells for each sample in each cell type cluster.

| **Subjects** |  |  | **Sample1** | **Sample2** | **Sample3** |
| --- | --- | --- | --- | --- | --- |
| **Number of total cells** |  |  | 5,663 | 7,147 | 13,521 |
| Cluster 6 | In_1 | Inhibitory<br>Neurons cells | 14.06% | 15.18% | 14.18% |
| Cluster 7 | In_6 |  |  |  |  |
| Cluster 0 | Ex_1 | Excitatory<br>Neurons cells | 72.63% | 75.68% | 68.12% |
| Cluster 1 | Ex_2 |  |  |  |  |
| Cluster 3 | Ex_4 |  |  |  |  |
| Cluster 2 | Ex_5 |  |  |  |  |
| Cluster 8 | Ex_6 |  |  |  |  |
| Cluster 4 | Ex_7 |  |  |  |  |
| Cluster 10 | Ex_8 |  |  |  |  |
| Cluster 9 | Astrocytes | Glia cells | 1.08% | 0.74% | 7.66% |
| Cluster 5 | Oligodendrocytes |  | 10.10% | 7.43% | 6.96% |
| Cluster 12 | OPC |  | 1.32% | 0.14% | 1.38% |
| Cluster 11 | Microglia | Non-neural<br>cells | 0.71% | 0.53% | 1.52% |
| Cluster 13 | Endothelial |  | 0.11% | 0.35% | 0.18% |

In inhibitory neuron, Ex excitatory neuron

**Figure 1:**
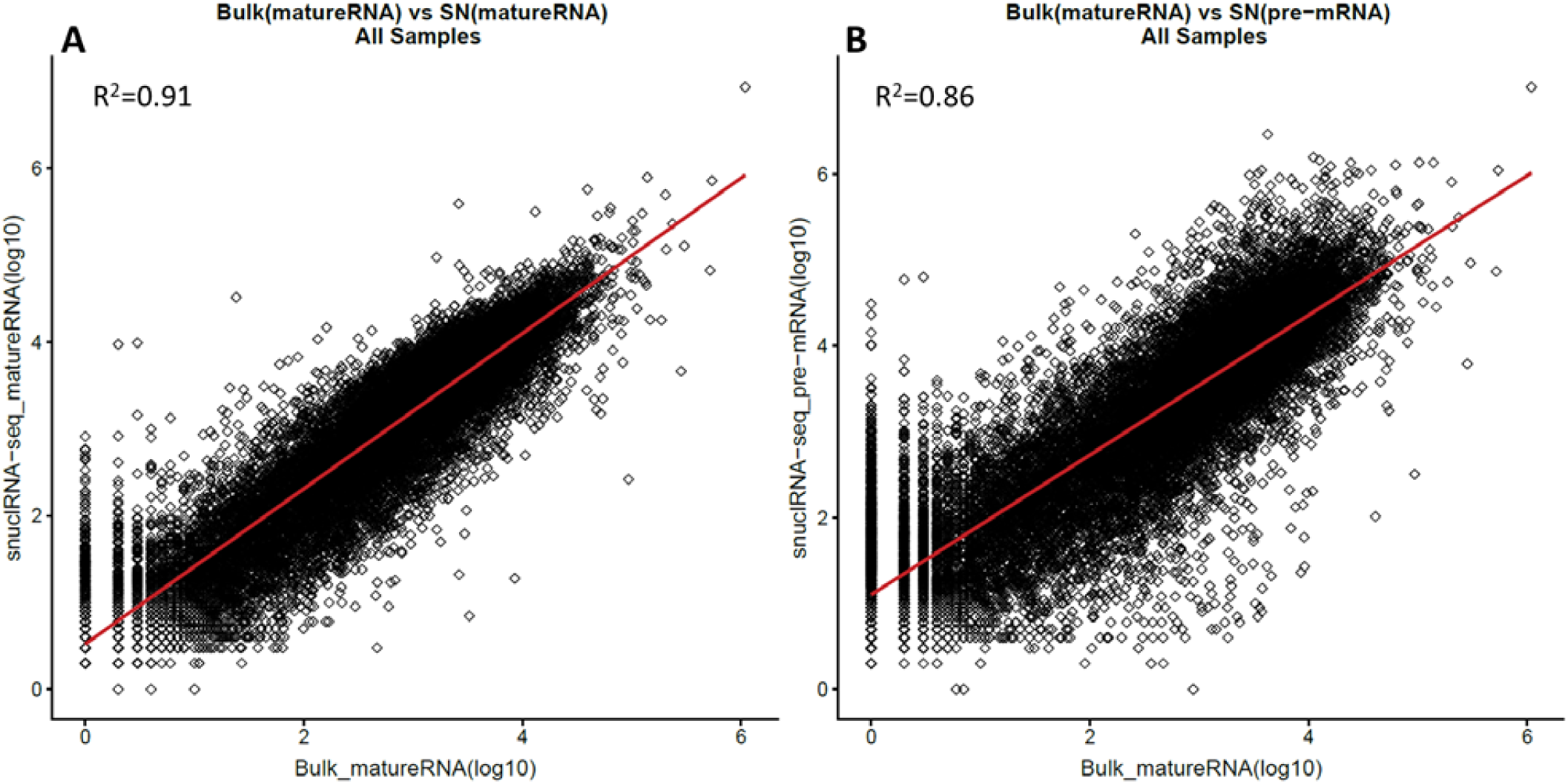
Correlation between Bulk-RNA-seq and Single Nuclei RNA-seq aligned using the pre-mRNA and mRNA annotation references. Along the X axis we show the gene expression values obtained from the bulk RNA-seq, and along the the Y-axis the single-nuclei expression, which was analyzed as bulk RNA-seq. (A) Bulk RNA-seq vs snuclRNA-seq aligned with mRNA (see Methods). (B) Bulk RNA-seq vs snuclRNA-seq aligned with pre-mRNA

These results indicated that the inclusion of introns for quantifying the gene expression of snuclRNA-seq data provides a better description of the nuclei specific expression profile, while conserving a correlation to the expression of bulk-RNA-seq, which validates the accuracy of this data.

### CGS approaches to cluster single nuclei from human postmortem brains

We initially sought to identify different cell types in brain samples by a CGS approach, that performs an unsupervised graph-based clustering [5] to identify groups of cells with similar expression profiles. This approach detected 25 clusters using 2,360 highly variable genes using the default resolution of 0.6, (Online Resource Table S4 and Figure 2-A). The clusters were annotated in six cell types (Figure 2-B, Online Resource Figure S2), however we noticed that many clusters included cells that were not evenly distributed among the three donors, but instead showed an overrepresentation or an underrepresentation of donors. Specifically, Sample2 is overrepresented in cluster 0, and underrepresented in clusters 2 and 3. Similarly, Sample3 is overrepresented in clusters 7 and 10 (Online Resource Table S4 and Figure 2-B). We employed Shannon’s Entropy to formally quantify the evenness of cell distribution among donors (Methods). This a metric that is maximized when the probability distribution is uniform. For the three brains we analyzed, it values ranged from 0 to 1.58. An even distribution of three brain samples in a cluster would result in entropy > 1.2. Lower values to this cutoff would be expected for a cluster with a 65% of overrepresentation of cells from a single donor or an underrepresentation of 10%. We observed that for clusters 0, 2, 3, 7, and 10 the entropy < 1.2 (Online Resource Table S4). Note that these clusters that exhibit uneven distribution of subjects included 9,781 nuclei, which represents 37.1% of the nuclei that passed QC. These clusters would not allow us to perform downstream comparisons on a relatively large proportion of the cells we sequenced, as many of these analyses require fair sampling of cells from the distinct donors.

**Figure 2:**
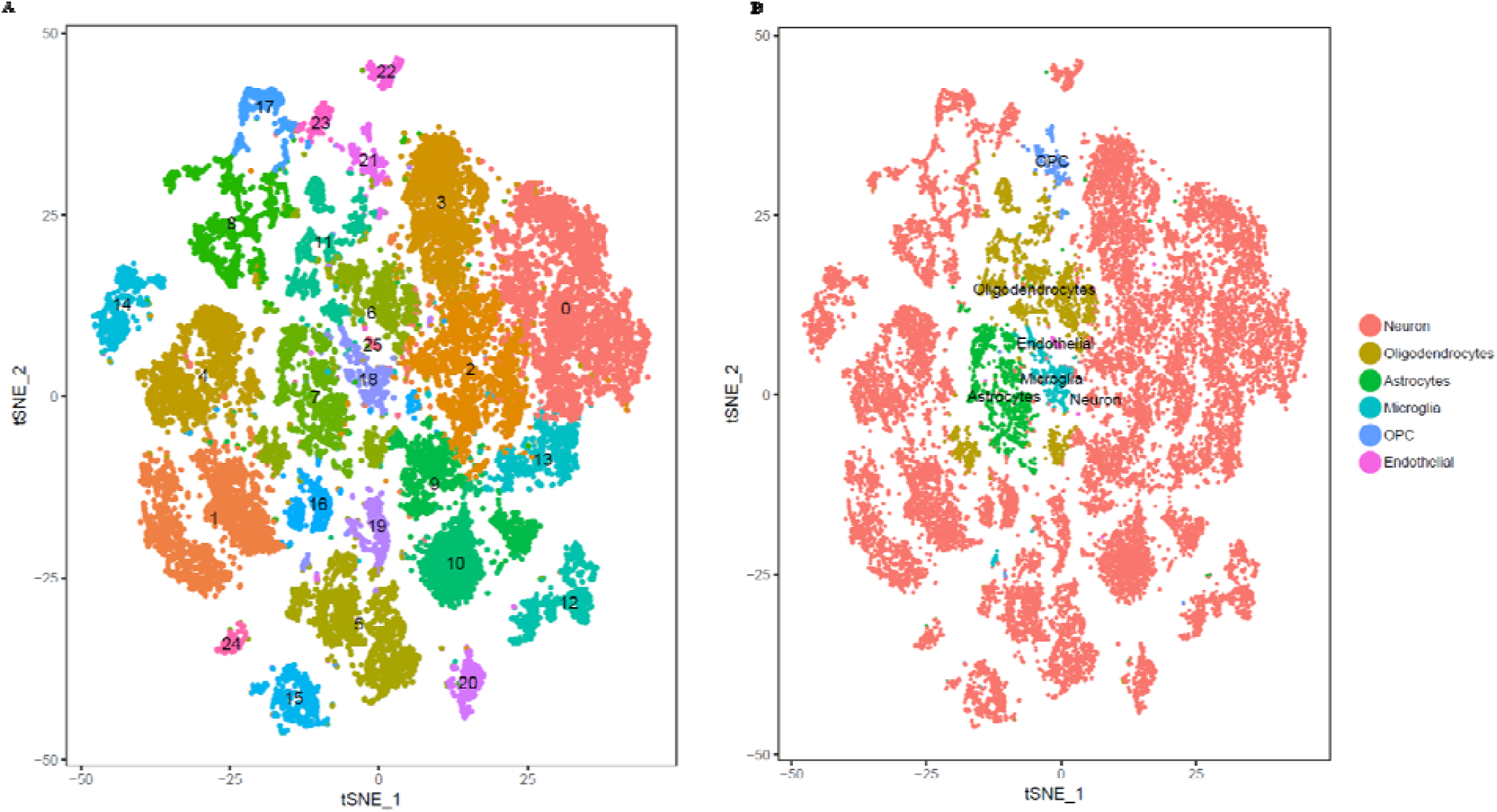
TSNE plots for the CGS Dimensional Reduction approach. TSNE plots depicting 26,331 nuclei.. Panel A) nuclei colored to represent the 25 CGS clusters. Panel B, Clusters annotated to represent cell types (Neuron, Ologodendrocytes, Astrocytes, Microglia, OPC and Endothelial).

We observed that generating a distinct number of clusters (by sampling the parameter resolution with values between 0.2 to 1.2) conserved the overall (nested) structure of the clusters. Higher resolution values tended to partition specific clusters, although cells were not reorganized in clusters with an even distribution of samples. When we employed lower resolution values (e.g. 0.2), the clusters started to show higher entropy values, as they grouped cells from all of the three donors. However, these clusters did not present cell type specific expression profiles. These results indicate that when analyzing snuclRNA-seq from human related brains, this approach produces some biases that constrain the comparisons for a large number of cells. This bias could be addressed by the reduction of the resolution in the analysis but the expression profile for these new clusters are not specific.

### The Hicat Gene Markers approach does not generate a trustworthy cell-type specific expression profile

Next, we evaluated whether clustering nuclei based on genes usually employed as cell type marker would produce cell type specific clusters. This approach, called “Hicat Gene Markers” [32], has been previously employed to analyze cell sorting (FACS) neurons. Using cell-type markers (Methods, Online Resources Table S1) we clustered nuclei into 23 bins (resolution 0.6 Online Resource Figure S3). We observed that the nuclei were better distributed among samples and the entropy values were closer to 1.58, (Online Resource Table S5) which is the maximum value that indicate perfectly even distribution. However, annotation of the clusters indicated that this method is not grouping homogenous cell types, as these clusters did not present cell type specific distinguishing expression profiles. Only two clusters were clearly annotated: Oligodendrocytes and Astrocytes (Online Resource Figure S3). These results suggest that expression of these markers, as captured by snuclRNA-seq, is not sufficient to cluster nuclei by cell type.

### The Consensus Gene Set (ConGen) generates clusters that show cell-type specific expression profiles

Finally, we envisioned an approach to generate data-driven clusters while avoiding biases introduced by overrepresentation and underrepresentation of donors in individual clusters. We explored whether the employment of common genes that are highly variable for each of the three donors would produce viable nuclei clusters (ConGen Methods 7.3). We detected 1,434 highly variable genes that we used to cluster nuclei and identified 14 bins (resolution 0.6). The nuclei in each of the clusters were evenly distributed among the three samples with overall high entropy values (Online Resource Table S6) and the expression profiles were specific enough to distinguish distinct cell types and subtypes (Online Resource Figure S6).

Based on these promising results, we generated gene expression imputed data to reduce the possible technical zeros (drop outs). From this new data, we identified 13 clusters (resolution 0.6, see Table 6). The gene expression patterns specific to cell type clusters were visualized using tSNE plot and DotPlot to represent the expression of gene markers of brain cell types (Figure 3 and Figure 4). A coincidence analysis identified a broad similarity between the clusters generated from the gene expression imputed data and the non-imputed data (Online Resource Figure S4). All of the clusters in the non-imputed data were also identified when we re-analyzed the imputed data, with the exception of two neuronal clusters (7 and 10) that were merged into a single cluster (cluster 6) after gene expression imputation.

**Figure 3:**
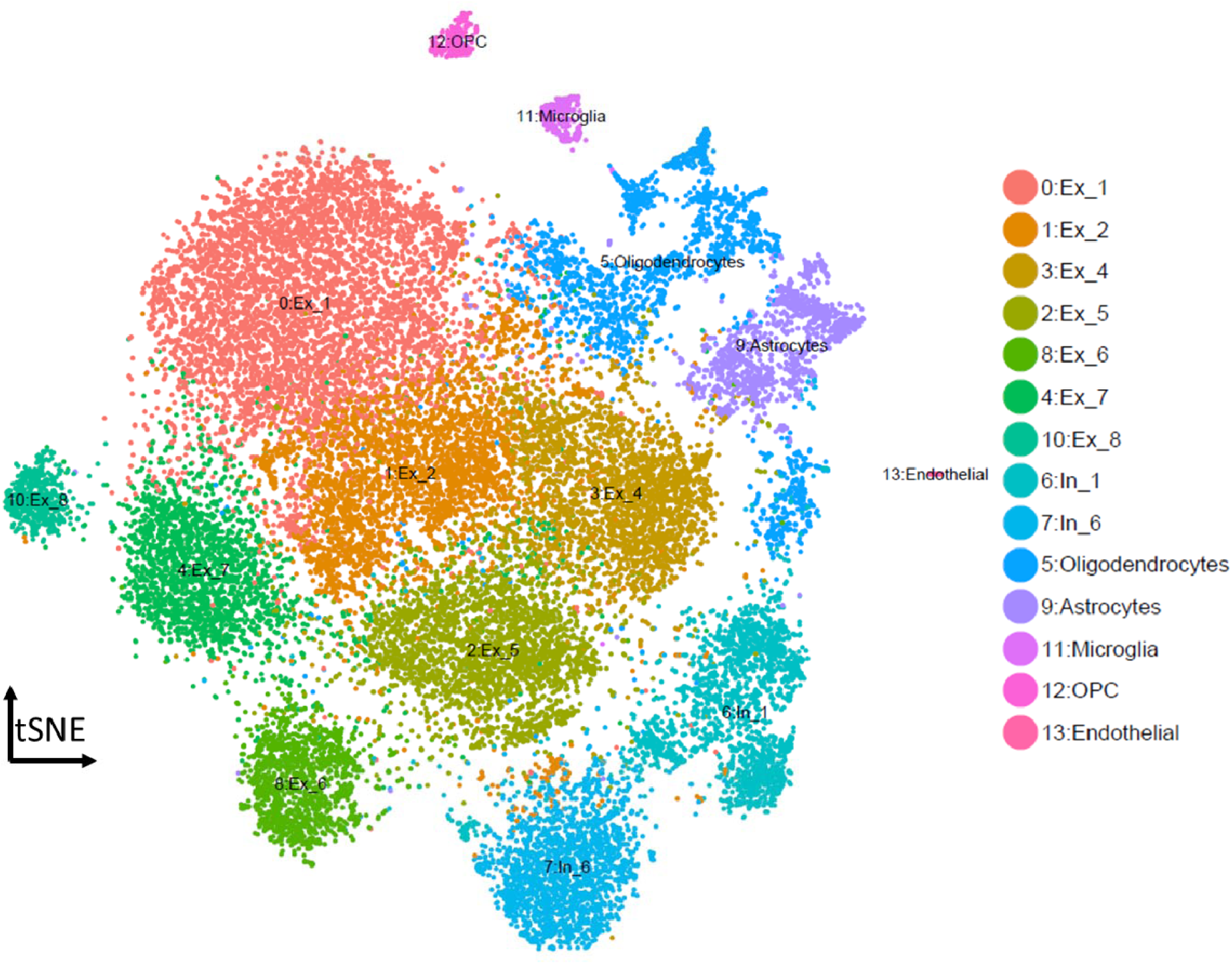
TSNE plots for Consensus Gene Set Dimensional Reduction approach. TSNE plots depicting 26,331 cells in 14 annotated clusters: Cluster0-Ex_1, Cluster1-Ex_2, Cluster3-Ex_4, Cluster2-Ex_5, Cluster8-Ex_6, Cluster4-Ex_7, Cluster10-Ex_8, Cluster6-In_1, Cluster7-In_6, Cluster5-Oligodendrocytes, Cluster9-Astrocytes, Cluster11-Microglia, Cluster12-OPC and Cluster 13-Endothelial. In inhibitory neuron, Ex excitatory neuron.

**Figure 4:**
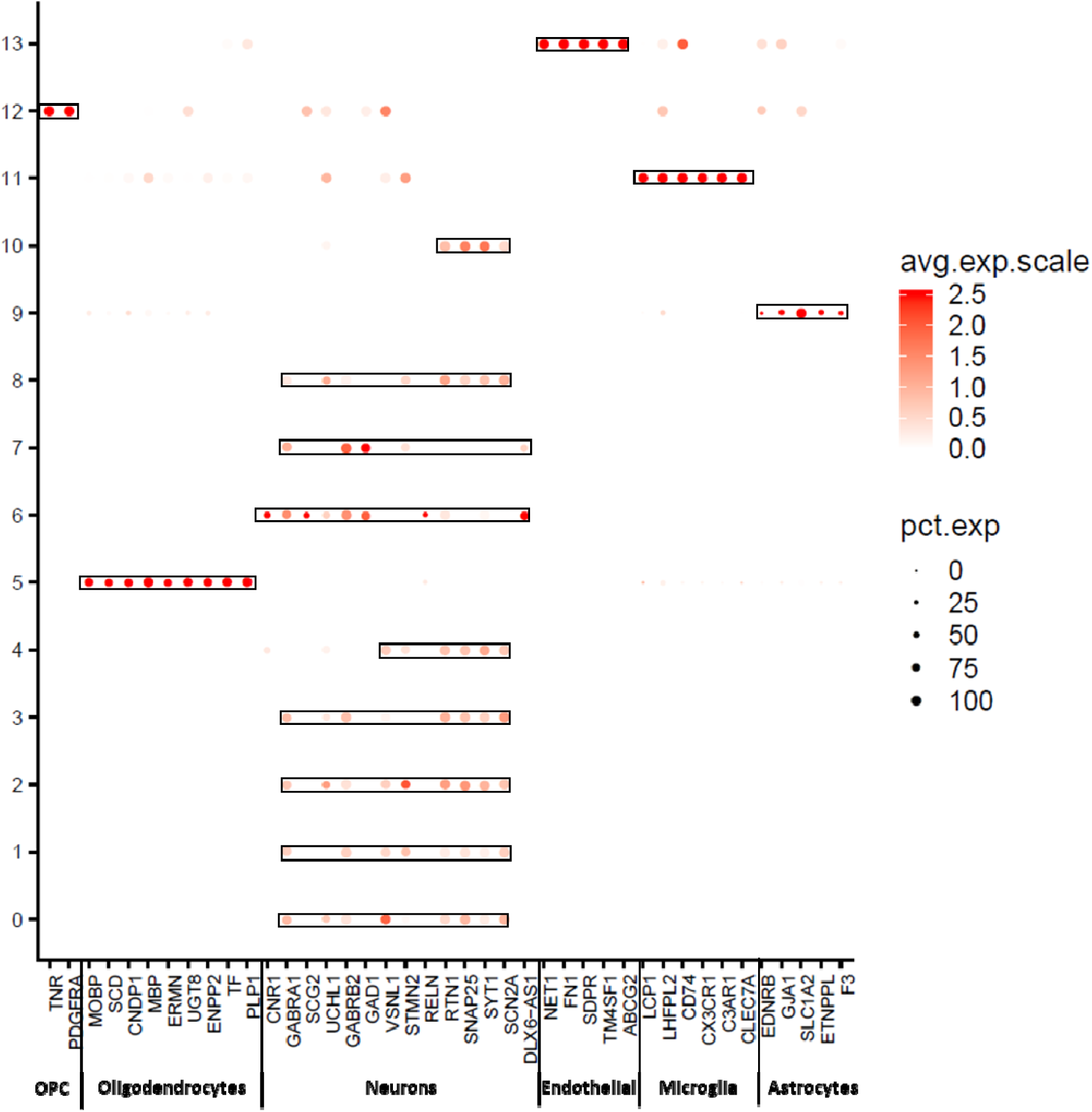
DotPlot depicting expression of markers genes selected by literature for the ConGen approach (see methods-Online Resource Table S2 and Table S3). This graphical approach allow us to annotated the clusters that were obtained by after the selection of the 1,434 common genes

Overall, based on the global expression patterns and similarity relationship among the clusters that we inferred (Methods), we were able to classify all the cells into three major groups, namely Neurons (both excitatory and inhibitory), glial cells (Astrocytes, Oligodendrocyte and OPC) and non-neural (Endothelial and Microglia) (Figure 5). Lake et al [20] established a series of neural sub markers that we employed to identify six subclasses of excitatory (Ex) neurons and 2 subclasses of inhibitory (In) neurons (Figure 6). The Inhibitory subgroups were distributed in different cerebral cortex layers (Figure 4). Inhibitory cell type 6 (In_6) were located in layer 2. The Inhibitory cell type 1 (In_1) were located between layer 2 to layer 5. In case of the excitatory neurons, we observed that most of the sub classes occupied multiple layers (Figure 4). In more detail, excitatory cell type 1 (Ex_1) covered the whole layer 2, while excitatory cell type 2 (Ex_2) from layer 2 to layer 4. Excitatory cell type 4 (Ex_4), excitatory cell type 5 (Ex_5) and excitatory cell type 6 (Ex_6) spread among layers 4 to 6. Finally, Excitatory cell type 8 (Ex_8) covered layers 5 to 6.

**Figure 5:**
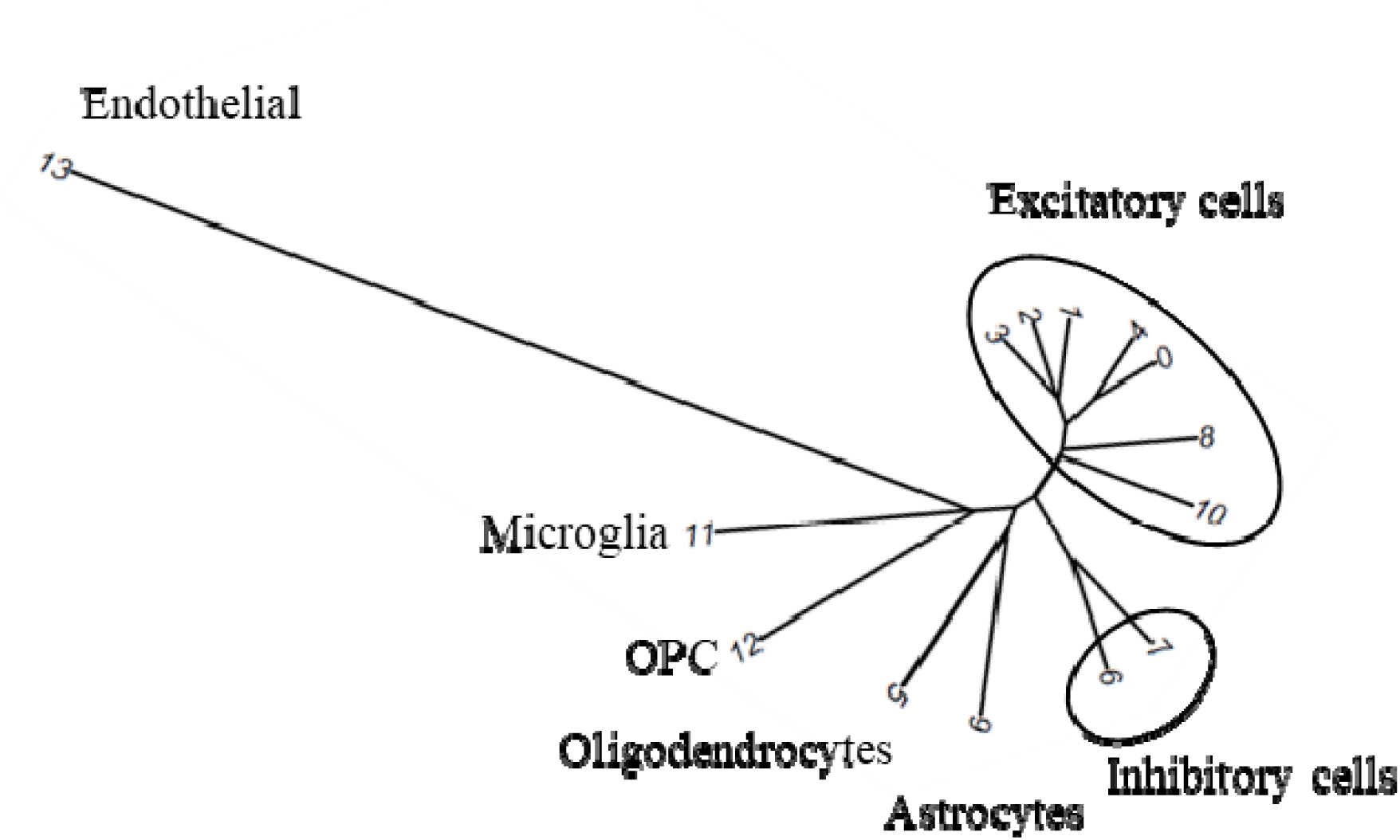
Dendrogram for Consensus Gene Set Clusters. This dendrogram shows the hierarchical relationship between clusters, based on the Euclidean distance of cluster mean expression. The proximity of that clusters 0, 1, 2, 3, 4, 8 and 10 indicates same cell type (Excitatory Neurons). Inhibitory neurons (clusters 6 and 7) are place in the same branch as excitatory neuron. This is another way to confirm the clustering obtaining by TSNE

**Figure 6:**
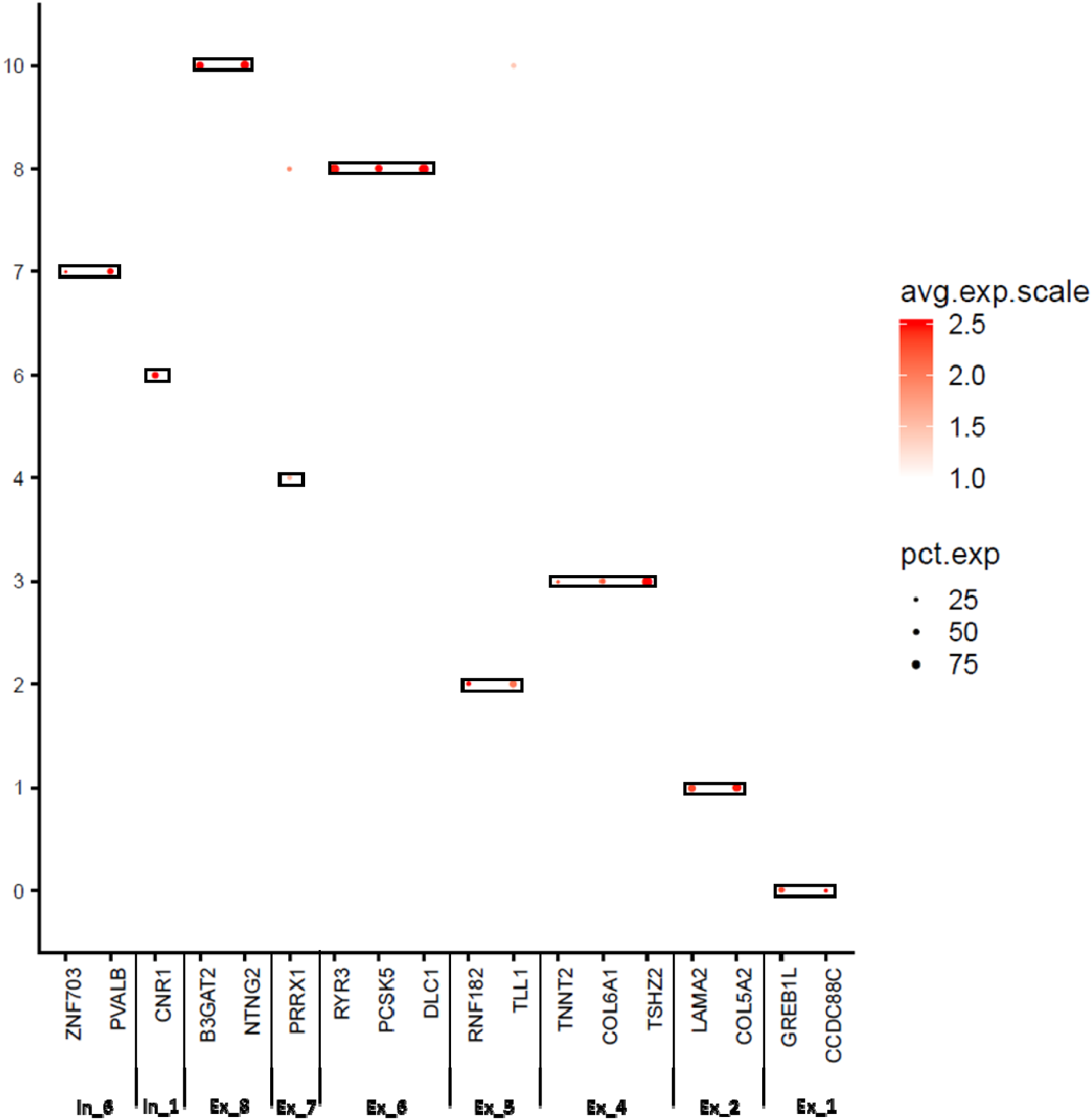
DotPlot depicting expression of the neurons cell for ConGen approach. Inhibitory neurons distributed between cluster 6 and 7, and the excitatory neurons distributed in cluster 0, 1, 2, 3, 4, 8 as defined by Lake et al [20]. In inhibitory neuron, Ex excitatory neuron

**Figure 7:**
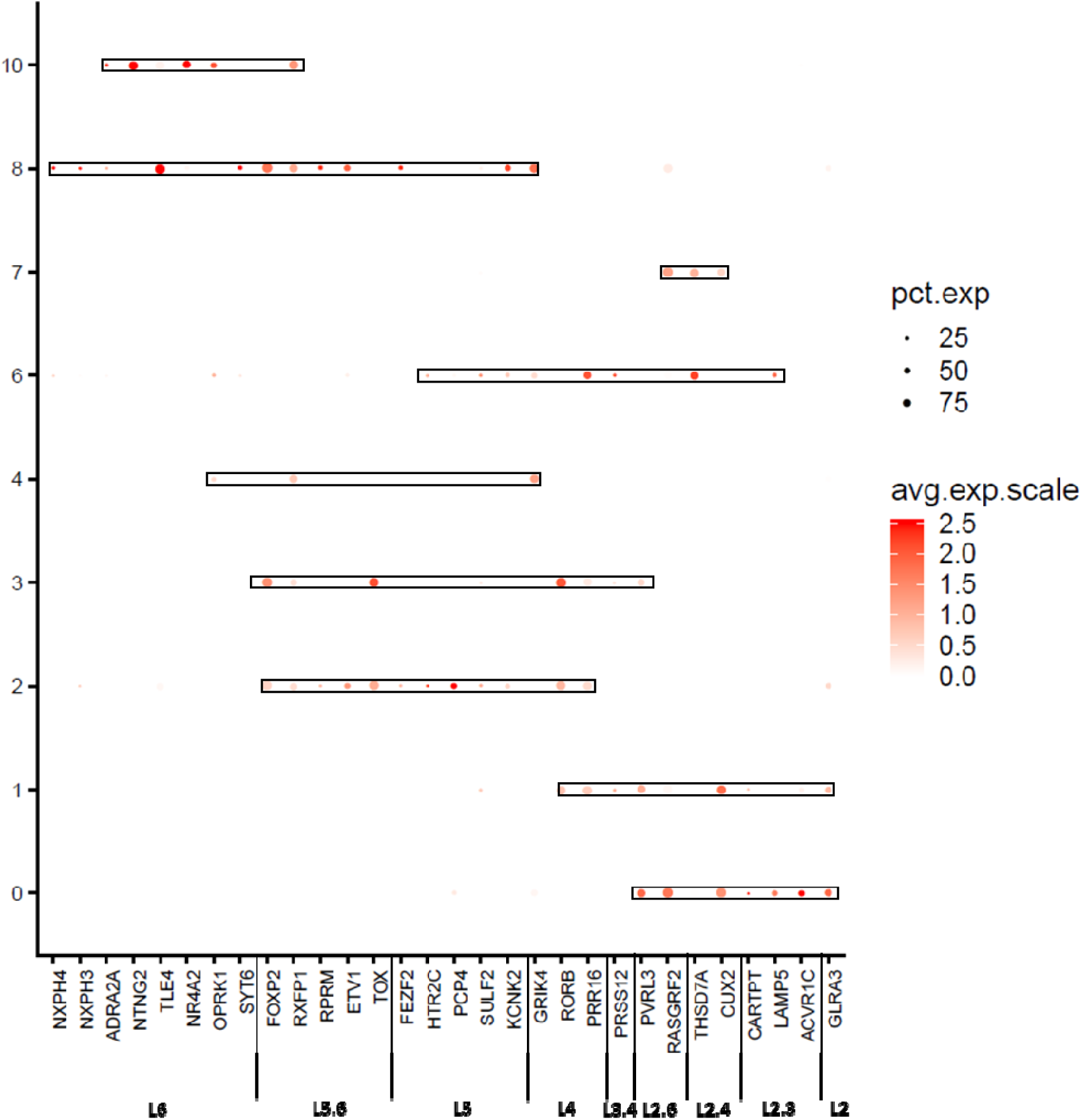
Dot Plots of Layer markers in different sub-clusters of Neuron cell sub-clusters from ConGen approach. DotPlot depicting expression of layers specific markers genes going from the superficial layers (e.g. L2) to the deeper layers (L6) for each Neural clusters (clusters 0, 1, 2, 3, 4, 6, 7, 8).

All of the clusters, except cluster 9 (Astrocytes), showed an entropy > 1.2 which indicates an even distribution of nuclei among the samples. Sample3 is overrepresented in cluster 9, as 90% of the cells are from this donor (Entropy = 0.89; Online Resource Figure S7), but no additional clusters included astrocytes for the other two samples. We evaluated whether our stringent QC process would remove astrocytes for Sample1 and Sample2 and introduce a bias in our data. To do so, we reduced the stringency of the QC parameters, and evaluated whether multi-modal distributions were modeling more accurately the number of genes expressed per nuclei, under the assumption that glial cells may express a lower number of genes [29], that would be removed during the cleaning process. However, using this multimodal distribution (Online Resources Figure S8) for QC and cleaning did not recover additional astrocytes nuclei. This suggest that some artifact prevented the efficient capture of astrocytes for these two samples. It has been previously reported that single-cell dissociation, capture, amplification and sequencing may distort brain cell abundances.[12, 29]. In particular, mouse brains staining were shown not to be correlated perfectly with scRNA-seq, suggesting that neurons were overrepresented relative to non-neurons [12, 29].

We performed coincidence analysis between the ConGen and CGS approaches (Online Resource Figure S6) to qualitatively compare these two approaches. Most of the neural cells from CGS were reorganized into neuronal clusters using the ConGen. This indicates that ConGen is not introducing any novel bias or artifact and nuclei are grouped by cell-type. Similarly, oligodendrocytes, which were grouped into two clusters by the CGS approach (clusters 6 and 11), were merged into a single cluster in the ConGen. The remaining cells were coincidently grouped into Endothelial, Oligodendrocyte precursor cell (OPC), Microglia and Astrocyte clusters by both approaches. Overall, these results indicated that the ConGen approach clusters cells in a manner that distinguishes cell type specific expression profiles that match those generated by the CGS approach, but reorganized neurons to avoid underrepresentation or overrepresentation of subjects.

### Single-nuclei RNA-seq reveals specific differences between the brain of the *PSEN1* p.A79V carrier and the two family members with sporadic AD

Neuronal cells accounted for 86.7%, 90.8% and 82.3% of the cells for Sample1, Sample2 and Sample3 respectively (Table 7). Once we broke down this number based on the neural cell subtype, we observed that the inhibitory neurons for Sample1, Sample 2 and Sample3 were 14.1%, 15.1% and 14.2%. However, the Sample3 (*PSEN1* carrier), showed fewer excitatory neurons (68.1%) compared to Sample1 (72.6%) and Sample2 (75.7%). Overall, this result supports our previous work [22], in which we reported a decreased neuronal percentage (20%) for carriers of pathological mutations in *APP, PSEN1* and *PSEN2*, and further suggests that this difference might be specific for excitatory neurons.

### Unsorted single-nuclei RNA-sequencing does not provide sufficient power to identify Disease Associated Microglia (DAM)

The analyses of sorted microglia cells from mouse models reported a novel microglia type associated with neurodegeneration [18], and the role of *TREM2* in their activation program [9]. This previous study performed immunohistochemical staining of human brain to identify the colocalization of Aβ particles with *SPP1*, a gene identified as a DAM marker [18]. Since then, much effort has been invested to identify the expression profile of DAM in human brains, including the analyses of bulk RNA-seq from homogenized human AD brains [9], and bulk RNA-seq from microglia sorted from ten fresh autopsy samples [25]. We hypothesize that the snuclRNA-seq data captures both the expression profiles of DAM cells proximal to Aβ plaques and microglia cells distal to Aβ plaques. We sought to investigate whether the expression profiles of microglia cells we called from the unsorted single-nuclei RNA-seq recapitulate the DAM marker genes. To do so, we performed an *in-silico* pseudo-time reconstruction to capture ordered sequence of the activation and transition to DAM cells from the transcriptomic profile of microglial cells [16]. From the 500 genes significantly associated with DAM [18] in mice AD models, we identified 326 homolog genes in the human genome. Single-nuclei RNA-seq detected the expression of 92 of these genes in microglia cells (Online Resources TablaS7). Surprisingly, 73 of these DAM markers show an expression significantly associated with the pseudo-time (q-value<0.05). To further investigate whether any artifact was introduced by the possibly distinct expression profiles of microglia from each donor, we performed subject specific analyses. We determined that only 20 DAM markers were significantly associated with pseudo-time when we analyzed the *PSEN1* carrier (Online Resources TablaS5). Similarly, 15 and 18 DAM markers were significantly associated with pseudo-time when we analyzed the microglial cells from the sporadic AD brains. Furthermore, five genes were consistently significantly associated in the microglia cells from all three samples, namely *EEF1A1, GLUL, KIAA1217, LDLRAD3, SPP1*. Curiously, *SPP1*, also referred as Osteopontin, is the only one of these genes among the top 10 DAM markers identified in mouse AD models, and previously employed in immunohistochemical staining of human brain [18]. Overall, these results suggest that unsorted single-nuclei RNA-seq from human brains does not provide enough microglia cells to clearly recapitulate initial findings of DAM marker genes, but instead promising results. We understand that sorting for glial cells and the development of novel methods that correct for subject specific biases would be optimally posed to provide more definitive results.

### A new web-based tool to explore the molecular atlas of Mendelian and Sporadic AD brains

Our web-based application (http://ngi.pub/snuclRNA-seq/) provides interactive access to the single-nuclei transcriptomic profiles of the three brains we analyzed. This application is designed to provide a user-friendly and comprehensive analysis of our data, through any modern web browser. The site features a graphical representation (tSEN projection) of the nuclei whose clusters are colored by their cell type, while allowing the selection of the genes with detected expression. For each gene, a graphical representation of its expression profile in the distinct cell types is produced, as well as a statistical interpretation of the differential expression among cell types.

We have regrouped all of the nuclei that were clustered in excitatory and inhibitory clusters that represented neuronal subtypes into a new cluster that represent all of the neurons (Figure 5). We have precomputed the statistical significance of the differential expression, and also the multiple comparisons correction p-value using Bonferroni multiple test correction. Using this tool, we validated the marker genes that we utilized to annotate the clusters (Table S2) are significantly over expressed for the expected cell types. Furthermore, this tool provides a single-nuclei cell-type specific expression reference that we believe will benefit additional research projects, including the annotation of data from other experiments, as well as helping to determine the expression profile of the distinct brain cell types of candidate genes identified in GWAS and sequencing projects.

## Discussion

The complexity and uniqueness of the cell types in the different regions and layers in the human brain make it difficult to understand the implication of each cell type in Alzheimer’s disease. Single-cell RNA-seq is a technology that is maturing rapidly. It is being used to generate a detailed molecular atlas of the brain and explore AD-induced damage at a cellular level. However, this technology is constrained to the analyses of fresh tissue. Single-nuclei RNA-sequencing (snuclRNA-seq) is still in its initial developmental stages. In this study, we showed that it is feasible to employ it to ascertain highly informative and unique brain tissue collected during many years and stored in brain banks. We showed that it is possible to ascertain a sufficient and diverse number of transcripts to identify different cell types from three frozen brains from family members with different forms of Alzheimer’s disease. To our knowledge, this is the first study to accomplish this.

Using this data, we generated a highly detailed molecular map of human brains for a carrier of a Mendelian mutation in *PSEN1* and two sporadic ADs. We profiled three major groups of cells in the brain, Neurons, glial cells (Astrocytes, Oligodendrocyte and OPC) and non-neuronal (Endothelial and Microglia). Furthermore, we identified distinct types of inhibitory and excitatory neurons. To do so, we extended current de facto methods to correctly process and analyze snuclRNA-seq data, and optimize the number of reads, genes and cells available for downstream analyses.

We showed the limitations and biases introduced by alternative approaches, CGS and Hicat Gene Markers, which were proven to produce reliable results for single-cell RNA-seq. The main problems that we faced were either the uneven representation of subjects in many clusters, or clusters with cells that do not show a cell-type specific expression profile. These problems originated from the approach employed to select the genes to cluster nuclei. Although these methods were effective to cluster cells from single-cell RNA-sequencing of fresh tissue or cloned animal models, they were not successful for snuclRNA-seq data

Our approach called ConGent is based on the identification of a common set of highly variable genes in common among the family related brains. This allowed us to infer clusters that represent distinct cell types and subtypes, while providing an approximately even representation of cells from all of the subjects in the clusters.

We used this information to analyze the cellular population structure to distinguish structures that are specific to Mendelian AD and differ from sporadic AD. Our results showed a reduced percentage of excitatory neurons in the brain carrier of a *PSEN1* mutation in comparison to two sporadic AD brains. This finding is concordant with our previous observations that Mendelian AD has distinctive and significantly decreased neuronal cell proportions. This result allows us to hypothesize that this phenomena is specific for excitatory neurons in carriers than sporadic AD, but additional samples should be study to verify this hypothesis.

We used the microglial cells to study their differential transcriptomic profiles. To our knowledge, this is the first attempt to study DAM markers using unsorted snuclRNA-seq from a *PSEN1 p.A79V* carrier and related AD brains. We showed that snuclRNA-seq provides accurate information, but we understand that the limited number of microglial cells sequenced from unsorted brain does not provide enough samples to perform unbiased analyses. In addition, we think that methods that identify genes associated with inferred pseudo-temporal ordering of the cells should correct for any donor specific effects that might confound the analyses. Still, our analysis replicated the association of Osteopontin (*SPI1*) with a temporal trajectory. We believe that by increasing the number of brains and by sorting the nuclei to capture glial cells, we will provide the power required to analyze and detect the expression profile and trajectories of DAM markers in AD brains.

Although all these results are encouraging, we recognize the limitation posed by the small sample size. In addition, we could not identify any family related neuropathology-free sample to employ as control. While under-powered, it is interesting to note that some of these results replicate the results of previous studies that were designed to provide sufficient statistical power to comparisons. Overall, we believe that this work proposes best practices for the generation, processing and analyses of single-nuclei RNA-seq (Figure 8) data that maximizes the amount of information able to be extracted from the samples. These are the lessons we learned while analyzing these brains: i) the quantification of nuclei using a “pre-mRNA” annotation will significantly increase the quantity of nuclei and the quantification of their expression profile. ii) Identification of genes that are highly variable in common among brain nuclei produce clusters with an even representation of all of the subjects. iii) Nuclei can be clustered differently using different resolution, but in general, they are assigned to clusters that are annotated to group nuclei from the same specific cell type. In addition, we identified a hierarchical relationship among the clusters as a product of different approaches or levels of resolution. To reveal these relationships, we propose coincidence analyses. iv) Shannon’s information theory of Entropy should be as a quantitative measure of even distribution of all samples in all clusters. v) To annotate each cluster, we need to use a consensus set of genes markers for each cell type from the current literature. There is not a standard set of gene markers. vi) A hierarchical clustering of clusters should reproduce expected results, grouping together neuronal subtypes in one branch and in another branch glial cells.

**Figure 8:**
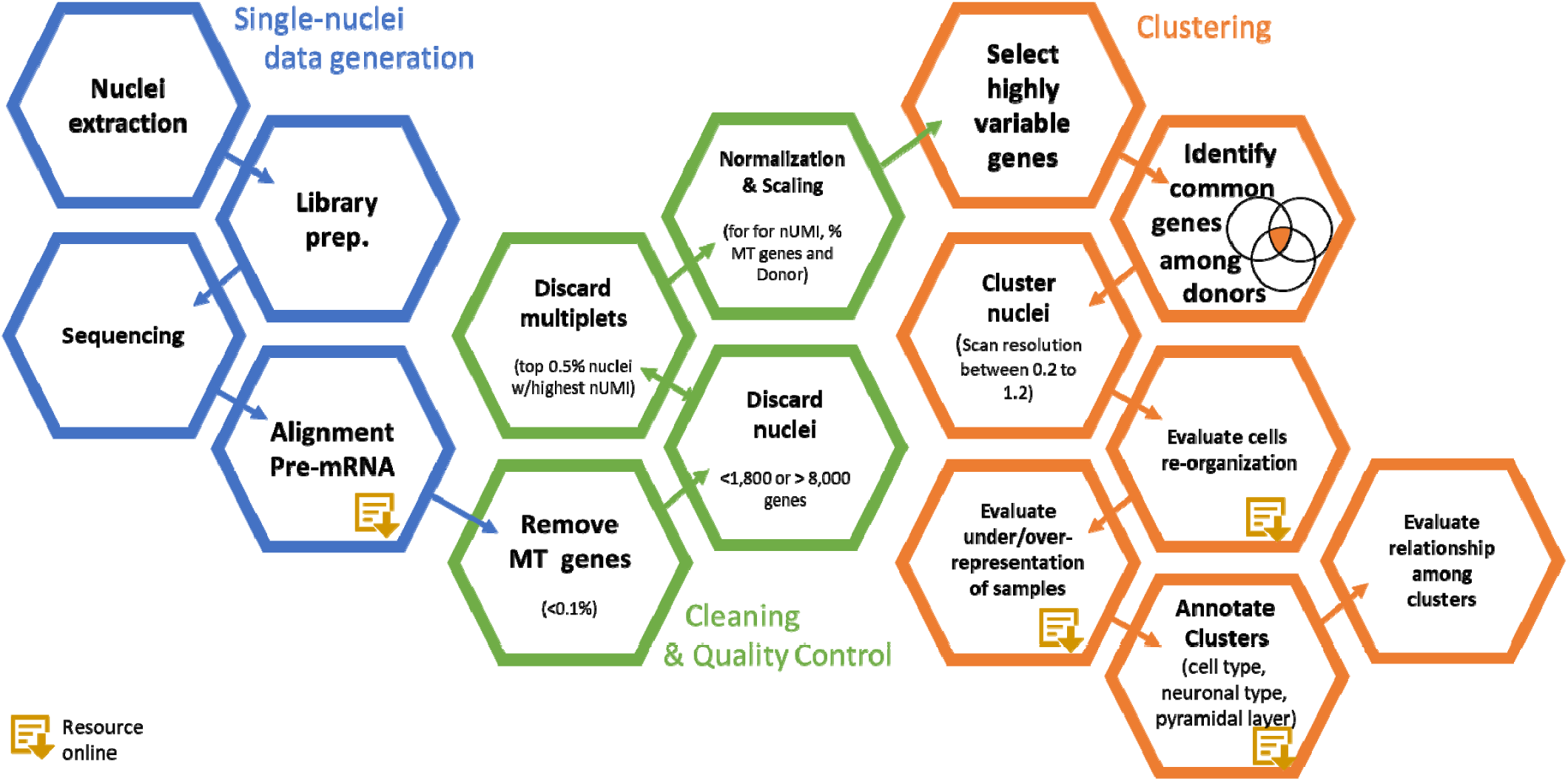
Workflow Analysis Plan. In blue, the single-nuclei data generation. The most important step is the quantification of nuclei using a “pre-mRNA” annotation. This step will significantly increase the quantity of nuclei and the quantification of their expression profile. In green, the cleaning and quality control steps. The QC followed standard measurements such as re moving mitochondrial genes (MT), removing doublets and multiples as well as the normalization of the data using nUMI, percent mitochondrial reads sample origin as confounding factors. In orange, the Clustering. In this step, we performed the identification of genes that are highly variable in common among brain nuclei from all of the subjects. Later on, the nuclei can be clustered differently using different resolution, but in general, they are assigned to clusters that are annotated to group nuclei from the same specific cell type. Next, we identified a hierarchical relationship among the clusters by performing coincidence analyses. The Entropy, from Shannon’s information theory, provides a quantitative measure of even distribution of samples in a cluster. To annotate the clusters, we use a set of genes markers for each cell type collected from the literature. Finally, a hierarchical clustering of clusters should reproduce expected results, grouping together neuronal subtypes in one branch and in another branch glial cells.

Finally, we have generated a highly detailed molecular atlas of AD brains that we are making available through an interactive, user-friendly, web-based application. We believe it will help not only the annotation of other snuclRNA-seq studies, but also additional high-throughput neurodegenerative genomic studies.

## Acknowledgments

We thank all participants and their families for their commitment and dedication to helping advance research into the early detection and causation of AD; and the Knight-ADRC research and support staff at each of the participating sites for their contributions to this study

## Author Contributions

JLDA analyzed the single-nuclei data and wrote the manuscript. ZL analyzed the bulk RNA-seq data. UD curated the neuropathological reports and phenotypic data. JPB perform genetic analyses to verify the identity of the samples, JD extracted the nuclei and JD and BAB collaborated with the manuscript edition. CC and OH designed and supervised the study and wrote the project. All authors read and approved the manuscript

## Compliance with ethical standards

Parietal lobe tissue of post-mortem brain and genetic information was obtained with informed consent for research use and were approved by Washington University in St. Louis Institutional Review Board.

## Funding

This work was supported by grants from the National Institutes of Health (R01AG057777, R01AG044546, P01AG003991, RF1AG053303, R01AG035083, and R01NS085419), the Alzheimer Association (NIRG-11-200110, BAND-14-338165, and BFG-15-362540). We thank the McDonnell Center for Cellular and Molecular Neurobiology for funds provided to generate the data we analyzed. BAB is supported by 2018 pilot funding from the Hope Center for Neurological Disorders and the Danforth Foundation Challenge at Washington University. The recruitment and clinical characterization of research participants at Washington University were supported by NIH P50 AG05681, P01 AG03991, and P01 AG026276.

We would like to thank the operations staff at the Elizabeth H. and James S. McDonnell III Genome Institute at Washington University for their assistance in constructing the single-nuclei RNA-seq libraries and generating sequence data for our project. This work was also supported by access to equipment made possible by the Hope Center for Neurological Disorders and the Departments of Neurology and Psychiatry at Washington University School of Medicine.

