## Supplementary Materials for "A single-nuclei RNA sequencing study of Mendelian and sporadic AD in the human brain"

* To whom correspondence should be addressed:

Carlos Cruchaga, PhD

Oscar Harari, PhD

Assistant Professor

Department of Psychiatry

Washington University School of Medicine

Campus Box 8134 | 425 S. Euclid Ave |BJC Institute of Health |Office: 9615 | St. Louis, MO 63110

Jorge L Del-Aguila:

Zeran Li:

Umber Dube:

Kathie A. Mihindukulasuriya:

John P Budde:

Maria Victoria Fernandez:

Laura Ibanez:

Joseph Bradley:

Fengxian Wang^:^

Kristy Bergmann:

Richard Davenport:

‎John C. Morris:

David M. Holtzman: ‎

Bruno A. Benitez:

Richard J. Perrin:

Joseph Dougherty:

Carlos Cruchaga:

Oscar Harari:

| **Table S1: 118 Cell type markers for Hicat Gene Marker** | |
| --- | --- |
| Cell-Type | Marker |
| Astrocytes | RYR3 |
|  | LRRC16A |
|  | GPR98 |
|  | F3 |
|  | ETNPPL |
|  | CLU |
|  | SLC1A2 |
|  | GJA1 |
|  | DIO2 |
|  | SLC4A4 |
|  | EDNRB |
|  | AQP4 |
|  | GPR37L1 |
|  | ALDOC |
|  | CPE |
|  | ATP1B2 |
|  | SLCO1C1 |
| Endotheial cells | DCN |
|  | RGS5 |
|  | ABCG2 |
|  | TM4SF1 |
|  | APOLD1 |
|  | SDPR |
|  | FN1 |
|  | NET1 |
| Microglia | B3GNT5 |
|  | CLEC7A |
|  | CCL4L1 |
|  | C3AR1 |
|  | CX3CR1 |
|  | CD74 |
|  | HLA-DRA |
|  | PLEK |
|  | LHFPL2 |
|  | HAVCR2 |
|  | LCP1 |
|  | CD53 |
|  | GPR183 |
|  | OLR1 |
|  | CH25H |
|  | IL1B |
|  | CD83 |
|  | MSR1 |
|  | CCL4 |
|  | C3 |
| Neurons | SYNPR |
|  | DLX6-AS1 |
|  | SCN2A |
|  | SYT1 |
|  | SNAP25 |
|  | RTN1 |
|  | RELN |
|  | STMN2 |
|  | VSNL1 |
|  | GAD1 |
|  | GABRB2 |
|  | UCHL1 |
|  | SCG2 |
|  | GABRA1 |
|  | CNR1 |
|  | PCLO |
| Oligodendrocytes | SEPT4 |
|  | PLP1 |
|  | TF |
|  | ENPP2 |
|  | SLC24A2 |
|  | TTLL7 |
|  | UGT8 |
|  | CLDND1 |
|  | ERMN |
|  | MBP |
|  | CNDP1 |
|  | SCD |
|  | MOBP |
|  | OPALIN |
|  | ANLN |
|  | SLAIN1 |
|  | CNP |
|  | SLC44A1 |
|  | SOX2-OT |
|  | TMEM144 |
|  | PIP4K2A |
|  | TULP4 |
|  | QDPR |
| OPC | PDGFRA |
|  | CNTN1 |
|  | TNR |
| Layer 2 | GLRA3 |
| Layer 2.3 | ACVR1C |
|  | LAMP5 |
|  | CARTPT |
| Layer 2.4 | CUX2 |
|  | THSD7A |
| Layer 2.6 | RASGRF2 |
|  | PVRL3 |
| Layer 3.4 | PRSS12 |
| Layer 4 | PRR16 |
| Layer 4.5 | RORB |
| Layer 4.6 | GRIK4 |
| Layer 5 | KCNK2 |
|  | SULF2 |
|  | PCP4 |
|  | HTR2C |
|  | FEZF2 |
| Layer 5.6 | TOX |
|  | ETV1 |
|  | RPRM |
|  | RXFP1 |
|  | FOXP2 |
| Layer 6 | SYT6 |
|  | OPRK1 |
|  | NR4A2 |
|  | TLE4 |
|  | NTNG2 |
|  | ADRA2A |
|  | NXPH3 |
|  | NXPH4 |

| **Table S2: Cell-type markers** | |  |  |
| --- | --- | --- | --- |
| Cell Type | Markers | Log Fold Change | Adjusted  p-value |
| Astrocytes | F3 | 0.40 | <2.23×10^-308^ |
|  | ETNPPL | 0.64 | <2.23×10^-308^ |
|  | SLC1A2 | 2.46 | <2.23×10^-308^ |
|  | GJA1 | 0.67 | <2.23×10^-308^ |
|  | EDNRB | 0.26 | 3.06×10^-122^ |
| Microglia | CLEC7A | 0.13 | <2.23×10^-308^ |
|  | C3AR1 | 0.09 | <2.23×10^-308^ |
|  | CX3CR1 | 0.57 | <2.23×10^-308^ |
|  | CD74 | 0.50 | <2.23×10^-308^ |
|  | LHFPL2 | 0.73 | 5.12×10^-178^ |
|  | LCP1 | 0.21 | <2.23×10^-308^ |
| Endothelial cells | ABCG2 | 2.11 | <2.23×10^-308^ |
|  | TM4SF1 | 1.11 | <2.23×10^-308^ |
|  | SDPR | 1.25 | <2.23×10^-308^ |
|  | FN1 | 1.38 | 1.44×10^-204^ |
|  | NET1 | 1.81 | 2.13×10^-132^ |
| Neurons | DLX6-AS1 | 0.10 | 5.48×10^-09^ |
|  | SCN2A | 0.53 | <2.23×10^-308^ |
|  | SYT1 | 0.79 | <2.23×10^-308^ |
|  | SNAP25 | 0.58 | <2.23×10^-308^ |
|  | RTN1 | 0.40 | <2.23×10^-308^ |
|  | STMN2 | 0.42 | 2.94×10^-240^ |
|  | VSNL1 | 0.48 | <2.23×10^-308^ |
|  | GAD1 | 0.16 | 1.60×10^-27^ |
|  | GABRB2 | 0.65 | <2.23×10^-308^ |
|  | UCHL1 | 0.24 | 2.21×10^-75^ |
|  | GABRA1 | 0.46 | <2.23×10^-308^ |
|  | CNR1 | 0.18 | 3.32×10^-05^ |
| Oligodendrocytes | PLP1 | 2.65 | <2.23×10^-308^ |
|  | TF | 1.95 | <2.23×10^-308^ |
|  | ENPP2 | 1.52 | <2.23×10^-308^ |
|  | UGT8 | 1.66 | <2.23×10^-308^ |
|  | ERMN | 1.19 | <2.23×10^-308^ |
|  | MBP | 2.19 | <2.23×10^-308^ |
|  | CNDP1 | 1.40 | <2.23×10^-308^ |
|  | SCD | 1.29 | <2.23×10^-308^ |
|  | MOBP | 1.76 | <2.23×10^-308^ |
| OPC | PDGFRA | 0.70 | <2.23×10^-308^ |
|  | TNR | 1.84 | 3.19×10^-167^ |

| **Table S3: Cerebral cortex layer-specific markers** | |
| --- | --- |
| **Layer type** | **Markers** |
| L2 | GLRA3 |
| L2-3 | ACVR1C |
|  | LAMP5 |
|  | CARTPT |
| L2-4 | CUX2 |
|  | THSD7A |
| L2-6 | RASGRF2 |
|  | PVRL3 |
| L3-4 | PRSS12 |
| L4 | PRR16 |
| L4-5 | RORB |
| L4-6 | GRIK4 |
| L5 | KCNK2 |
|  | SULF2 |
|  | PCP4 |
|  | HTR2C |
|  | FEZF2 |
| L5-6 | TOX |
|  | ETV1 |
|  | RPRM |
|  | RXFP1 |
|  | FOXP2 |
| L6 | SYT6 |
|  | OPRK1 |
|  | NR4A2 |
|  | TLE4 |
|  | NTNG2 |
|  | ADRA2A |
|  | NXPH3 |
|  | NXPH4 |

| **Table S4: Distribution of nuclei per subject using CGS** | | | | |
| --- | --- | --- | --- | --- |
| **Subject** | **Sample1** | **Sample2** | **Sample3** | **Entropy*** |
| **Number of total cells** | 5,663 | 7,147 | 13,521 |  |
| Cluster 0 | 18.33% | 34.14% | 0.77% | 1.03 |
| Cluster 1 | 8.79% | 8.87% | 8.68% | 1.58 |
| Cluster 2 | 10.21% | 0.34% | 10.09% | 1.10 |
| Cluster 3 | 0.37% | 0.01% | 13.31% | 0.19 |
| Cluster 4 | 7.33% | 7.16% | 6.58% | 1.58 |
| Cluster 5 | 6.11% | 6.90% | 7.09% | 1.58 |
| Cluster 6 | 3.00% | 4.60% | 6.07% | 1.53 |
| Cluster 7 | 0.81% | 0.77% | 8.88% | 0.76 |
| Cluster 8 | 4.10% | 4.28% | 5.35% | 1.57 |
| Cluster 9 | 7.82% | 3.47% | 3.85% | 1.48 |
| Cluster 10 | 0.09% | 0.00% | 8.17% | 0.11 |
| Cluster 11 | 7.28% | 3.43% | 1.90% | 1.38 |
| Cluster 12 | 3.67% | 4.14% | 1.92% | 1.52 |
| Cluster 13 | 6.04% | 5.11% | 0.24% | 1.12 |
| Cluster 14 | 2.74% | 2.85% | 1.82% | 1.56 |
| Cluster 15 | 3.21% | 1.65% | 2.12% | 1.53 |
| Cluster 16 | 1.24% | 1.30% | 2.86% | 1.47 |
| Cluster 17 | 2.38% | 2.21% | 1.37% | 1.55 |
| Cluster 18 | 0.85% | 0.52% | 2.73% | 1.24 |
| Cluster 19 | 1.13% | 2.63% | 1.24% | 1.47 |
| Cluster 20 | 0.78% | 1.85% | 1.25% | 1.50 |
| Cluster 21 | 1.59% | 0.31% | 1.62% | 1.34 |
| Cluster 22 | 0.67% | 1.48% | 0.53% | 1.44 |
| Cluster 23 | 0.72% | 1.01% | 0.72% | 1.57 |
| Cluster 24 | 0.62% | 0.62% | 0.67% | 1.58 |
| Cluster 25 | 0.12% | 0.35% | 0.18% | 1.45 |
| *Entropy values <1.2 indicates uneven sample representation in the cluster | | | | |

| **Table S5: Distribution of nuclei per subject using Hicat Marker Gene** | | | | |
| --- | --- | --- | --- | --- |
| **Subject** | **Sample1** | **Sample2** | **Sample3** | **Entropy*** |
| **Number of total cells** | 5,663 | 7,147 | 13,521 |  |
| Cluster 0 | 24.28% | 22.16% | 21.93% | 1.58 |
| Cluster 1 | 16.07% | 11.49% | 23.06% | 1.53 |
| Cluster 2 | 5.60% | 8.74% | 4.53% | 1.53 |
| Cluster 3 | 3.36% | 2.00% | 4.36% | 1.52 |
| Cluster 4 | 3.62% | 6.28% | 1.90% | 1.43 |
| Cluster 5 | 4.29% | 4.14% | 2.57% | 1.55 |
| Cluster 6 | 2.15% | 5.97% | 2.06% | 1.39 |
| Cluster 7 | 2.45% | 2.15% | 3.14% | 1.57 |
| Cluster 8 | 2.38% | 2.46% | 2.98% | 1.58 |
| Cluster 9 | 3.30% | 1.94% | 2.89% | 1.55 |
| Cluster 10 | 3.67% | 2.74% | 2.25% | 1.56 |
| Cluster 11 | 2.91% | 1.37% | 3.22% | 1.50 |
| Cluster 12 | 2.15% | 3.20% | 2.54% | 1.57 |
| Cluster 13 | 2.40% | 5.67% | 1.11% | 1.30 |
| Cluster 14 | 2.97% | 1.55% | 2.96% | 1.53 |
| Cluster 15 | 1.73% | 2.90% | 2.70% | 1.55 |
| Cluster 16 | 2.91% | 2.63% | 2.27% | 1.58 |
| Cluster 17 | 1.40% | 2.29% | 2.89% | 1.53 |
| Cluster 18 | 2.03% | 4.03% | 1.24% | 1.42 |
| Cluster 19 | 1.77% | 1.37% | 2.71% | 1.53 |
| Cluster 20 | 3.67% | 2.14% | 1.27% | 1.46 |
| Cluster 21 | 1.38% | 0.55% | 2.80% | 1.33 |
| Cluster 22 | 1.96% | 1.43% | 1.32% | 1.56 |
| Cluster 23 | 1.54% | 0.77% | 1.30% | 1.53 |
| *Entropy values <1.2 indicates uneven sample representation in the cluster | | | | |

| **Table S6: Distribution of nuclei per subject using Consensus Gene Set** | | | | |
| --- | --- | --- | --- | --- |
| **Subjects** | **Sample1** | **Sample2** | **Sample3** | **Entropy*** |
| **Number of Total cells** | 5,663 | 7,147 | 13,521 |  |
| Cluster 0 | 16.95% | 27.05% | 18.96% | 1.55 |
| Cluster 1 | 18.67% | 16.97% | 9.74% | 1.53 |
| Cluster 2 | 9.25% | 11.82% | 10.61% | 1.58 |
| Cluster 3 | 9.01% | 3.85% | 14.41% | 1.41 |
| Cluster 4 | 9.43% | 7.61% | 8.82% | 1.58 |
| Cluster 5 | 10.01% | 7.33% | 6.74% | 1.56 |
| Cluster 6 | 6.55% | 7.18% | 7.70% | 1.58 |
| Cluster 7 | 4.82% | 5.61% | 5.09% | 1.58 |
| Cluster 8 | 6.20% | 6.87% | 3.76% | 1.54 |
| Cluster 9 | 1.09% | 0.71% | 7.54% | 0.90 |
| Cluster 10 | 2.93% | 2.49% | 1.60% | 1.54 |
| Cluster 11 | 2.98% | 1.51% | 1.97% | 1.53 |
| Cluster 12 | 0.67% | 0.50% | 1.50% | 1.42 |
| Cluster 13 | 1.32% | 0.14% | 1.38% | 1.23 |
| Cluster 14 | 0.11% | 0.35% | 0.18% | 1.42 |
| *Entropy values <1.2 indicates uneven sample representation in the cluster. | | | | |

| **Table S7 DAM markers significantly associated with pseudo-time** | | | | | | | | |
| --- | --- | --- | --- | --- | --- | --- | --- | --- |
| **DAM** | **pval** | **qval** | **pval_Sample1** | **qval_Sample1** | **pval_Sample2** | **qval_Sample2** | **pval_Sample3** | **qval_Sample3** |
| KIAA1217 | 0 | 0 | 1.7E-104 | 1.9E-102 | 6.5E-09 | 2.6E-07 | 6.4E-87 | 6.4E-85 |
| GLUL | 1.9E-39 | 6.5E-38 | 1.8E-05 | 0.00028 | 4.07E-17 | 2.8E-15 | 1.0E-09 | 1.9E-08 |
| SAMD4A | 4.7E-34 | 1.4E-32 | 3.9E-07 | 8.3E-06 | 0.045 | 0.182 | 1.1E-28 | 4.4E-27 |
| CD74 | 1.5E-31 | 4.1E-30 | 0.000 | 0.0001 | 1.4E-13 | 8.1E-12 | 0.194 | 0.255 |
| SCD | 8.1E-29 | 2.1E-27 | 7.7E-08 | 1.7E-06 | 0.005 | 0.0456 | 0.188 | 0.249 |
| EEF1A1 | 3.6E-25 | 8.5E-24 | 0.000 | 0.003 | 8.5E-07 | 2.6E-05 | 0.010 | 0.033 |
| RFTN1 | 5.7E-24 | 1.3E-22 | 6.4E-16 | 2.9E-14 | 0.025 | 0.130 | 1.7E-20 | 6.2E-19 |
| COX4I1 | 2.3E-22 | 4.7E-21 | 0.025 | 0.095 | 5.7E-05 | 0.001 | 8.6E-08 | 1.3E-06 |
| MYO1E | 1.9E-20 | 3.9E-19 | 1.7E-08 | 4.1E-07 | 0.009 | 0.071 | 3.4E-09 | 6.1E-08 |
| SPP1 | 3.0E-20 | 5.9E-19 | 0.0004 | 0.005 | 0.003 | 0.029 | 4.1E-43 | 2.0E-41 |
| PGAM1 | 4.7E-19 | 8.3E-18 | 0.042 | 0.134 | 2.4E-06 | 6.7E-05 | 0.019 | 0.052 |
| B2M | 6.2E-18 | 1.1E-16 | 0.063 | 0.171 | 0.0002 | 0.004 | 0.021 | 0.057 |
| RHOB | 1.5E-17 | 2.4E-16 | 0.027 | 0.100 | 3.3E-07 | 1.1E-05 | 0.00013 | 0.001 |
| LDLRAD3 | 5.5E-17 | 8.6E-16 | 2.3E-07 | 4.9E-06 | 0.001 | 0.013 | 3.9E-12 | 9.0E-11 |
| PEA15 | 1.0E-16 | 1.5E-15 | 0.031 | 0.111 | 2.6E-06 | 7.3E-05 | 1.4E-12 | 3.4E-11 |
| MAMDC2 | 5.6E-13 | 5.9E-12 | 2.3E-10 | 6.9E-09 | 0.148 | 0.339 | 5.8E-13 | 1.4E-11 |
| CREG1 | 1.0E-12 | 1.0E-11 | 0.211 | 0.339 | 0.004 | 0.038 | 0.021 | 0.055 |
| TPI1 | 1.9E-12 | 1.9E-11 | 0.042 | 0.134 | 0.009 | 0.070 | 0.008 | 0.027 |
| RPL5 | 9.6E-11 | 7.8E-10 | 0.151 | 0.280 | 0.001 | 0.013 | 0.049 | 0.100 |
| RPS14 | 1.1E-10 | 8.9E-10 | 0.030 | 0.107 | 0.014 | 0.091 | 0.030 | 0.073 |
| GHR | 1.4E-10 | 1.1E-09 | 0.000 | 0.000 | 0.014 | 0.091 | 0.000 | 0.000 |
| RPL3 | 8.3E-10 | 5.7E-09 | 0.003 | 0.022 | 0.139 | 0.329 | 0.031 | 0.073 |
| TTYH2 | 5.7E-09 | 3.5E-08 | 0.006 | 0.035 | 0.005 | 0.043 | 0.250 | 0.307 |
| GPR34 | 8.4E-09 | 5.1E-08 | 0.042 | 0.134 | 0.013 | 0.088 | 0.122 | 0.186 |
| RPL11 | 1.2E-08 | 7.4E-08 | 0.028 | 0.102 | 0.107 | 0.290 | 0.178 | 0.240 |
| TMEM163 | 2.1E-08 | 1.2E-07 | 0.000 | 0.001 | 0.574 | 0.697 | 0.000 | 0.000 |
| RPS4X | 2.4E-08 | 1.3E-07 | 0.177 | 0.307 | 0.020 | 0.115 | 0.572 | 0.598 |
| SCPEP1 | 3.7E-08 | 2.0E-07 | 0.002 | 0.017 | 0.039 | 0.169 | 0.246 | 0.304 |
| RPS25 | 7.9E-08 | 4.1E-07 | 0.004 | 0.025 | 0.076 | 0.240 | 0.052 | 0.104 |
| RPS8 | 2.4E-07 | 1.1E-06 | 0.003 | 0.020 | 0.262 | 0.470 | 0.197 | 0.258 |
| PPT1 | 5.3E-07 | 2.3E-06 | 0.110 | 0.234 | 0.686 | 0.776 | 0.138 | 0.202 |
| SERPINE2 | 6.4E-07 | 2.2E-06 | 0.100 | 0.222 | 0.003 | 0.028 | 0.040 | 0.088 |
| DPP7 | 1.1E-06 | 4.4E-06 | 0.031 | 0.111 | 0.110 | 0.294 | 0.045 | 0.096 |
| ATP6V0E1 | 1.5E-06 | 5.8E-06 | 0.060 | 0.167 | 0.003 | 0.029 | 0.091 | 0.152 |
| CXCL14 | 1.8E-06 | 6.7E-06 | 0.228 | 0.355 | 0.648 | 0.750 | 0.551 | 0.579 |
| RNASET2 | 5.4E-06 | 1.9E-05 | 0.228 | 0.355 | 0.002 | 0.021 | 0.355 | 0.402 |
| FAU | 7.2E-06 | 2.4E-05 | 0.136 | 0.264 | 0.235 | 0.442 | 0.092 | 0.152 |
| LDHA | 9.3E-06 | 3.0E-05 | 0.115 | 0.238 | 0.039 | 0.169 | 0.073 | 0.131 |
| CD164 | 1.2E-05 | 3.8E-05 | 0.084 | 0.201 | 0.028 | 0.139 | 0.100 | 0.163 |
| RPLP1 | 1.3E-05 | 4.1E-05 | 0.077 | 0.191 | 0.393 | 0.570 | 0.158 | 0.221 |
| APOE | 1.5E-05 | 4.5E-05 | 0.003 | 0.024 | 0.202 | 0.405 | 0.033 | 0.077 |
| PRDX1 | 1.6E-05 | 4.8E-05 | 0.201 | 0.329 | 0.071 | 0.231 | 0.153 | 0.216 |
| CD83 | 1.9E-05 | 5.6E-05 | 0.002 | 0.018 | 0.522 | 0.660 | 0.404 | 0.445 |
| RPL37 | 3.3E-05 | 9.3E-05 | 0.132 | 0.259 | 0.057 | 0.206 | 0.019 | 0.052 |
| SDF4 | 4.1E-05 | 0.0001 | 0.054 | 0.157 | 0.207 | 0.411 | 0.157 | 0.220 |
| NRP1 | 6.4E-05 | 0.0002 | 0.456 | 0.549 | 0.090 | 0.265 | 1.6E-06 | 1.8E-05 |
| HINT1 | 7.4E-05 | 0.0002 | 0.265 | 0.389 | 0.073 | 0.235 | 0.185 | 0.247 |
| HLA-A | 0.0001 | 0.0003 | 0.383 | 0.488 | 0.654 | 0.753 | 0.396 | 0.439 |
| EGLN3 | 0.0002 | 0.0004 | 0.046 | 0.142 | 0.339 | 0.528 | 0.002 | 0.011 |
| BHLHE40 | 0.0002 | 0.0005 | 0.198 | 0.326 | 0.030 | 0.146 | 0.270 | 0.325 |
| RPLP0 | 0.0003 | 0.0006 | 0.408 | 0.509 | 0.406 | 0.580 | 0.339 | 0.388 |
| LGMN | 0.0003 | 0.0007 | 0.051 | 0.152 | 0.752 | 0.822 | 0.157 | 0.220 |
| EEF1B2 | 0.0004 | 0.0009 | 0.506 | 0.586 | 0.014 | 0.091 | 0.660 | 0.677 |
| BRI3BP | 0.0005 | 0.0011 | 0.225 | 0.352 | 0.164 | 0.362 | 0.009 | 0.030 |
| CTSA | 0.0006 | 0.0013 | 0.069 | 0.179 | 0.048 | 0.190 | 0.128 | 0.191 |
| PLBD2 | 0.0010 | 0.0020 | 0.183 | 0.314 | 0.137 | 0.327 | 0.030 | 0.073 |
| TOX2 | 0.0014 | 0.0025 | 0.287 | 0.408 | 0.723 | 0.800 | 0.190 | 0.252 |
| RAMP1 | 0.0014 | 0.0026 | 0.416 | 0.514 | 0.308 | 0.508 | 0.001 | 0.007 |
| RPL18 | 0.0016 | 0.0028 | 0.258 | 0.383 | 0.173 | 0.374 | 0.770 | 0.777 |
| RPS3 | 0.0027 | 0.0045 | 0.053 | 0.156 | 0.612 | 0.725 | 0.031 | 0.074 |
| RPL14 | 0.0032 | 0.0052 | 0.202 | 0.330 | 0.447 | 0.607 | 0.226 | 0.286 |
| RPS19 | 0.0035 | 0.0056 | 0.062 | 0.170 | 0.215 | 0.420 | 0.421 | 0.461 |
| P2RY13 | 0.0038 | 0.0060 | 0.800 | 0.828 | 0.303 | 0.504 | 0.643 | 0.662 |
| ITGAX | 0.0040 | 0.0063 | 0.298 | 0.419 | 0.116 | 0.302 | 0.100 | 0.163 |
| ANXA5 | 0.0092 | 0.0130 | 0.069 | 0.179 | 0.537 | 0.670 | 0.145 | 0.208 |
| RPL22 | 0.0106 | 0.0147 | 0.400 | 0.503 | 0.119 | 0.306 | 0.148 | 0.211 |
| RPL7 | 0.0125 | 0.0170 | 0.526 | 0.601 | 0.058 | 0.208 | 0.280 | 0.335 |
| SHISA5 | 0.0135 | 0.0183 | 0.887 | 0.904 | 0.286 | 0.492 | 0.088 | 0.148 |
| RPL37A | 0.0140 | 0.0188 | 0.040 | 0.130 | 0.680 | 0.772 | 0.004 | 0.016 |
| CSF2RA | 0.0141 | 0.0190 | 0.512 | 0.590 | 0.957 | 0.969 | 0.621 | 0.642 |
| SH3PXD2B | 0.0142 | 0.0190 | 0.071 | 0.181 | 0.399 | 0.574 | 0.052 | 0.104 |
| RPLP2 | 0.0176 | 0.0230 | 0.243 | 0.369 | 0.099 | 0.278 | 0.385 | 0.429 |
| RPL4 | 0.0226 | 0.0286 | 0.166 | 0.295 | 0.816 | 0.865 | 0.073 | 0.130 |
| IGF1 | 0.0252 | 0.0315 | 0.317 | 0.432 | 0.213 | 0.418 | 0.423 | 0.463 |
| RPS28 | 0.0265 | 0.0330 | 0.494 | 0.578 | 0.476 | 0.628 | 0.230 | 0.290 |
| MAF | 0.0297 | 0.0364 | 0.282 | 0.403 | 0.502 | 0.647 | 0.734 | 0.744 |
| DPYS | 0.0340 | 0.0411 | 0.068 | 0.178 | 0.581 | 0.703 | 0.446 | 0.484 |
| SULF2 | 0.0371 | 0.0444 | 0.113 | 0.237 | 0.757 | 0.825 | 0.115 | 0.179 |
| PTCHD1 | 0.0422 | 0.0495 | 0.370 | 0.476 | 0.523 | 0.661 | 0.019 | 0.052 |
| IFI44 | 0.0570 | 0.0647 | 0.629 | 0.684 | 0.727 | 0.803 | 0.465 | 0.500 |
| RPL10A | 0.0608 | 0.0686 | 0.284 | 0.405 | 0.556 | 0.685 | 0.284 | 0.338 |
| PLEKHH2 | 0.0623 | 0.0701 | 0.016 | 0.070 | 0.651 | 0.752 | 0.608 | 0.630 |
| CREB3L2 | 0.0627 | 0.0705 | 0.158 | 0.286 | 0.462 | 0.618 | 0.164 | 0.227 |
| CD84 | 0.0635 | 0.0713 | 0.293 | 0.414 | 0.388 | 0.567 | 0.802 | 0.807 |
| NPM1 | 0.0945 | 0.1031 | 0.259 | 0.383 | 0.668 | 0.763 | 0.523 | 0.554 |
| CTSS | 0.1104 | 0.1189 | 0.320 | 0.436 | 0.706 | 0.788 | 0.538 | 0.568 |
| DHRS3 | 0.1124 | 0.1210 | 0.854 | 0.875 | 0.620 | 0.730 | 0.205 | 0.265 |
| TXNIP | 0.1282 | 0.1370 | 0.075 | 0.189 | 0.477 | 0.628 | 0.573 | 0.598 |
| DKK2 | 0.3387 | 0.3465 | 0.474 | 0.561 | 0.705 | 0.788 | 0.383 | 0.428 |
| GAS2L3 | 0.4079 | 0.4146 | 0.085 | 0.202 | 0.856 | 0.895 | 0.585 | 0.608 |
| FAM20C | 0.4477 | 0.4537 | 0.507 | 0.586 | 0.606 | 0.719 | 0.380 | 0.425 |
| FLT1 | 0.8162 | 0.8187 | 0.855 | 0.875 | 0.753 | 0.823 | 0.277 | 0.332 |

**Figures**

**Figure S1**: Single Nuclei after extraction from sample1.

**Figure S2**: DotPlot depicting expression of marker genes selected from the literature (see methods-Online Resource Table S2 and Table S3) on Classic Gene Set data.

**Figure S3**: **Hicat Gene Markers Dimensional Reduction approach**. . Panel A) TSNE plots depicting 26,331 nuclei in 23 clusters and Panel B) DotPlot depicting expression of markers genes selected by literature (see methods-Online Resource Table S2 and Table S3).

**Figure S4**: **DotPlot depicting expression of marker genes.** Marker genes were selected by literature (see methods-Online Resource Table S2 and Table S3) on Consensus Gene Set data prior gene expression imputation

**Figure S5**: **Coincidence Analysis between Consensus Gene Set and its gene expression imputed counterpart**. This plot summarizes how the nuclei are (dis)similarly clustered by the two approaches. All of the clusters learned by the Consensus Gene Set approach are represented along the X axis, and along the Y axis the clusters from the imputed gene expression. The Jaccard index for each pair is represented by the size of the circle (the bigger the higher value), and the color represents the pointwise mutual information (blue: low values; red: high values). All of the clusters in the non-imputed data were also identified when we re-analyzed the imputed data, with the exception of two neuronal clusters (7 and 10) that were merged into a single cluster (cluster 6).

**Figure S6:** **Coincidence Analysis between Classic Gene Set and its Consensus Gene Set**. This plot summarizes how the nuclei are (dis)similarly clustered by the two approaches. All of the clusters learned by the Consensus Gene Set approach are represented along the X axis, and along the Y axis the clusters from the imputed gene expression. The Jaccard index for each pair is represented by the size of the circle (the bigger the higher value), and the color represents the pointwise mutual information (blue: low values; red: high values). Both approaches identified clusters similarly for Endothelial cells, OPC, Microglia and Astrocytes. Neural cells from CGS approach were reorganized into clusters also annotated as neuronal by the Consensus Gene Set approach (shown in boxes).

**Figure S7**: **Entropy Data**. Graphical representation of Entropy values for the three cell clusters methodologies used. On the y-axis are nuclei counts, on x-axis are Entropy values from 0 to 1.58. Sample 3 is overrepresented in cluster 9 for the ConGen as well as in cluster 7 for the CGS in both cases these clusters were annotated as Astrocytes.

**Figure S8**: Multimodal distributions of unique molecular identified (UMI) per nuclei.

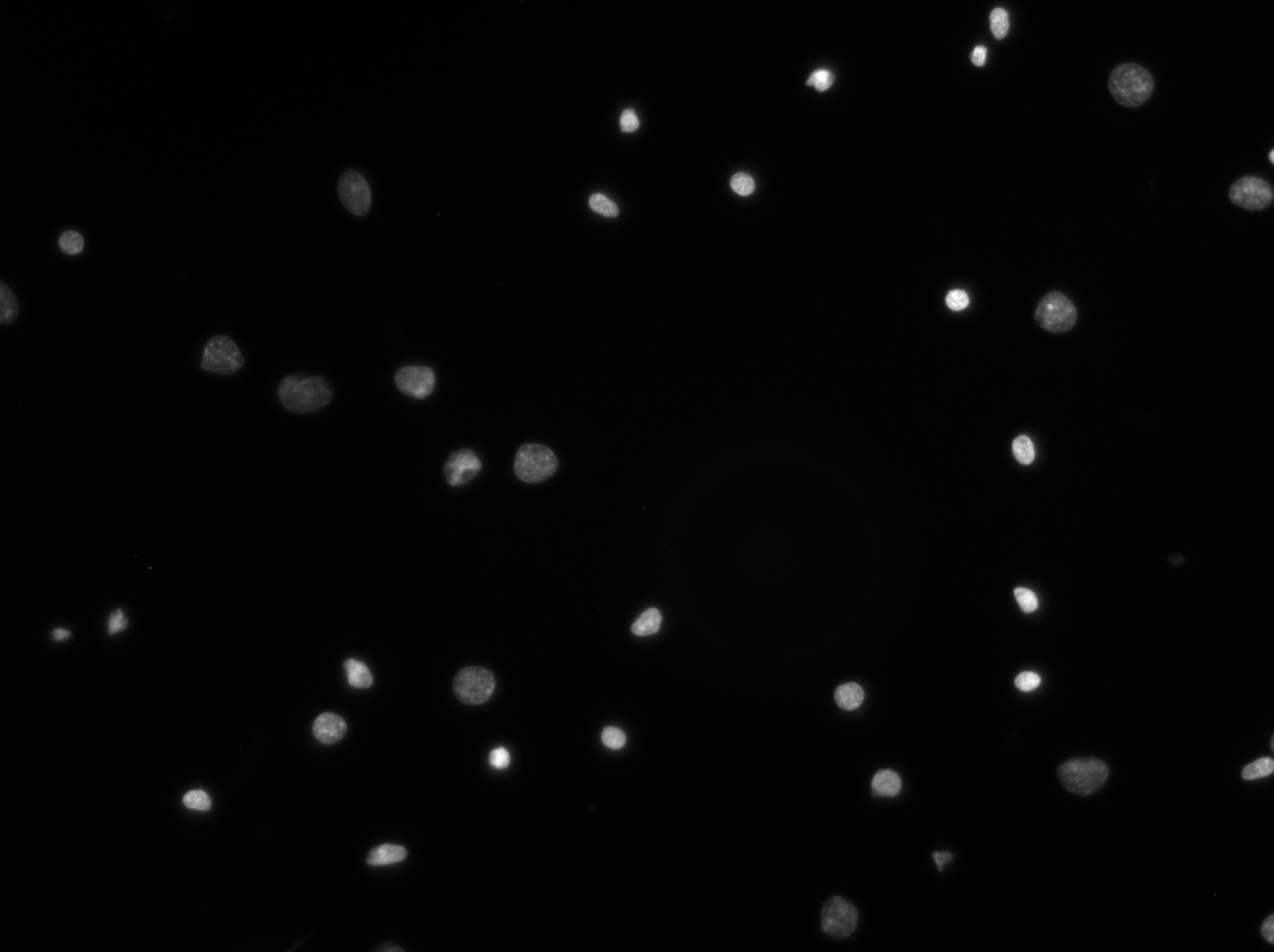

**Figure S1**

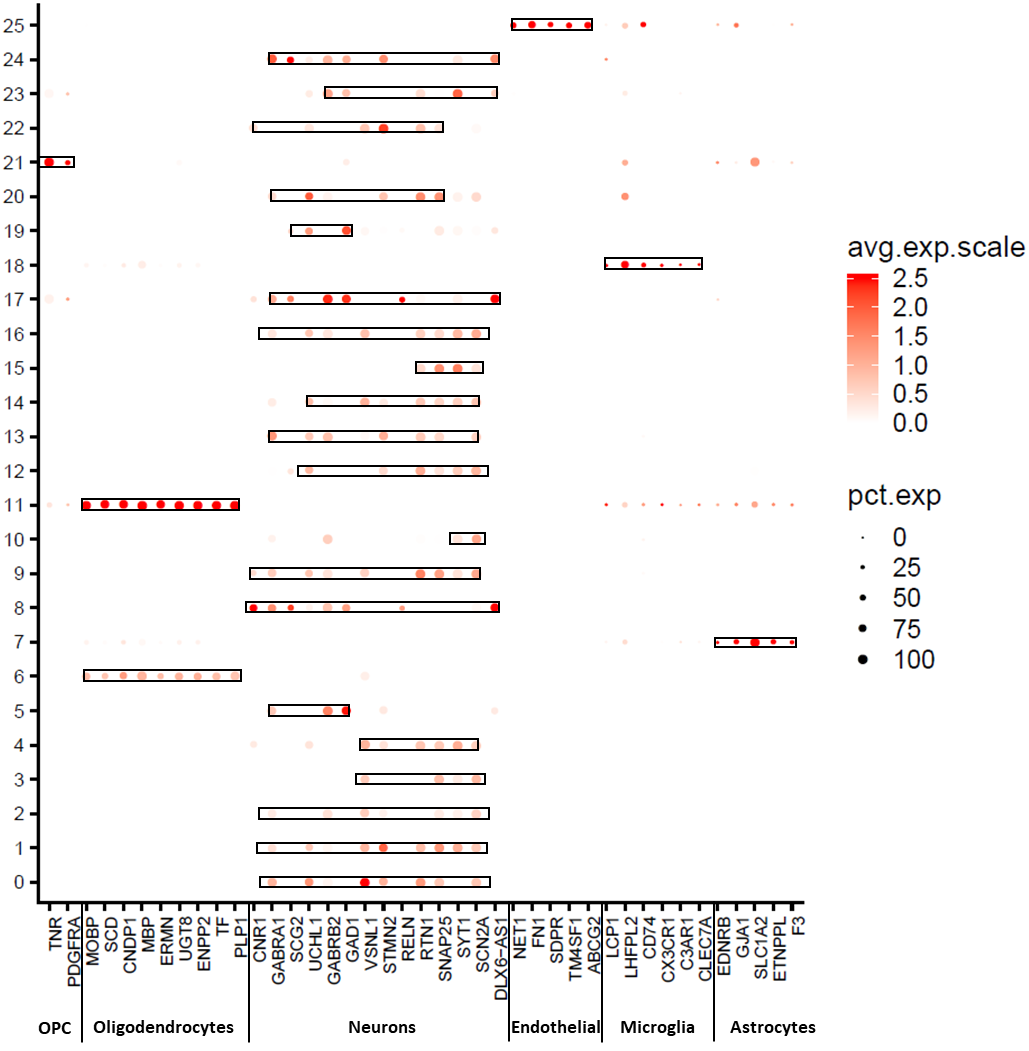

**Figure S2**

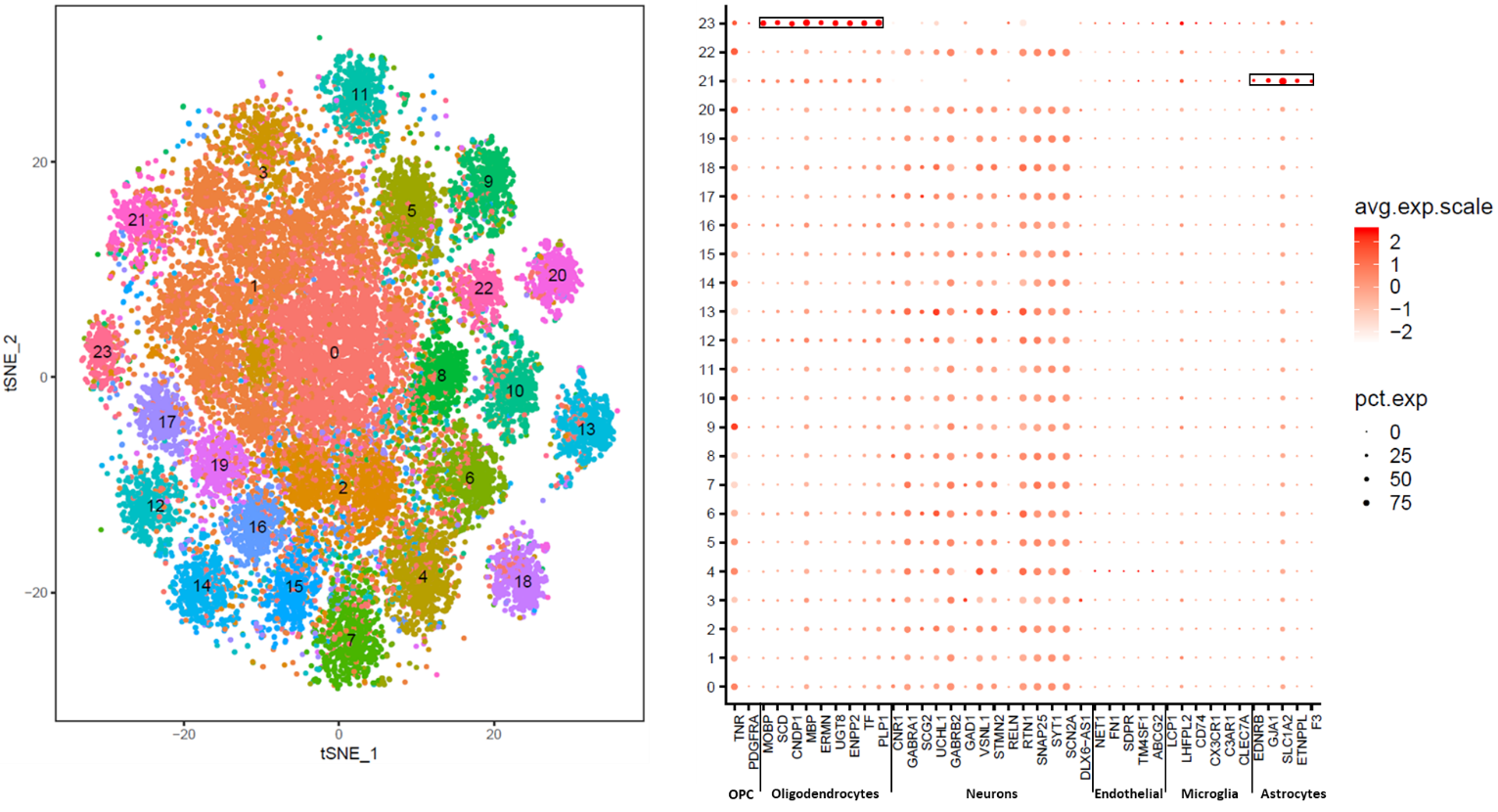

**Figure S3**

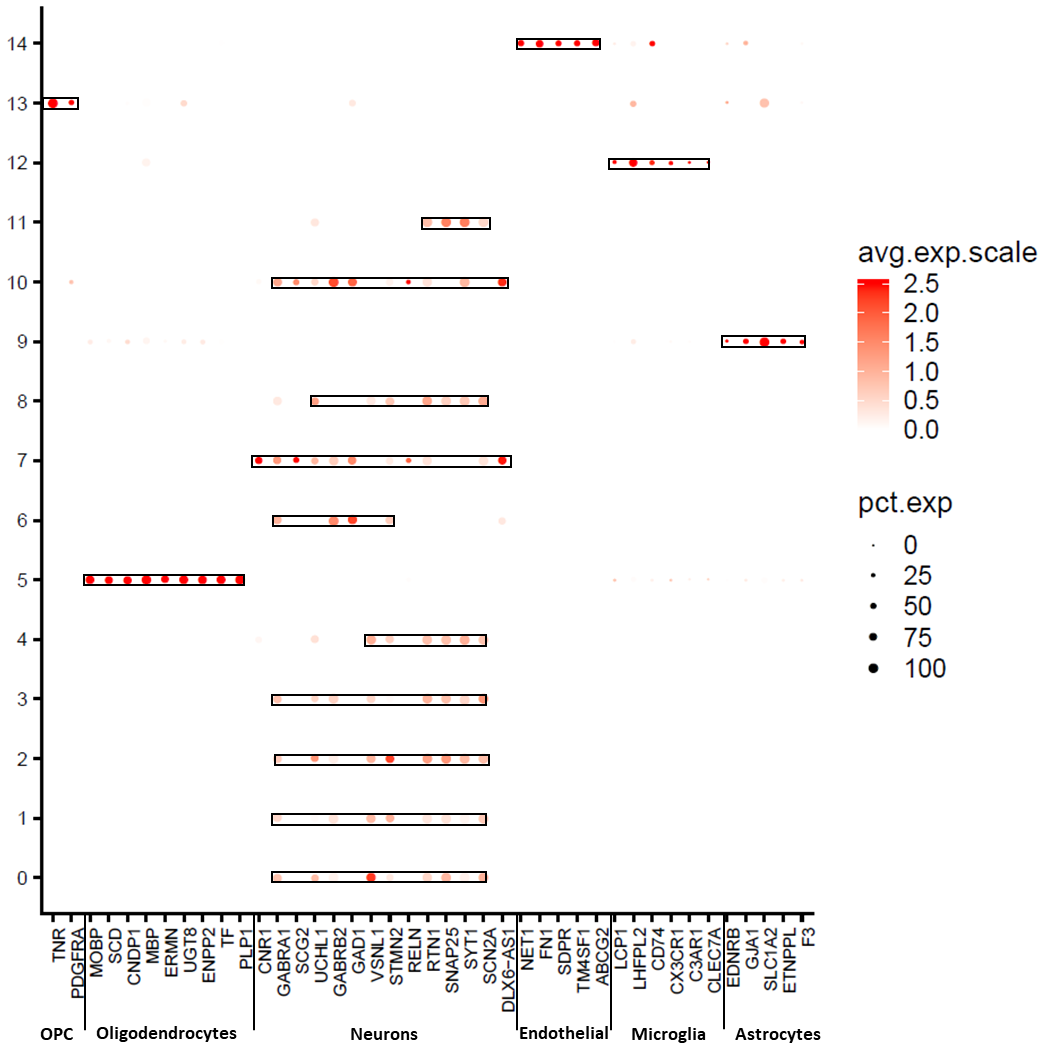

**Figure S4**

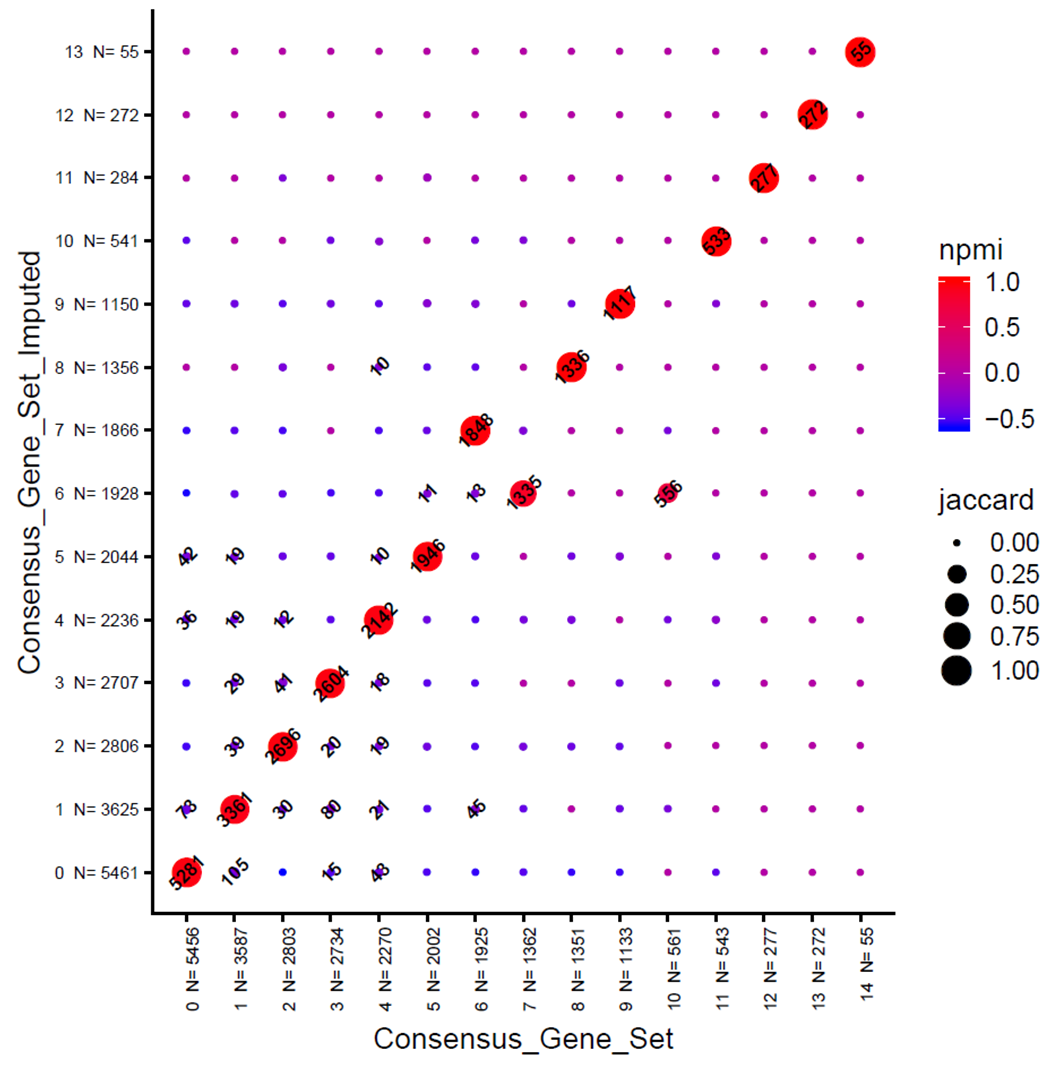

**Figure S5**

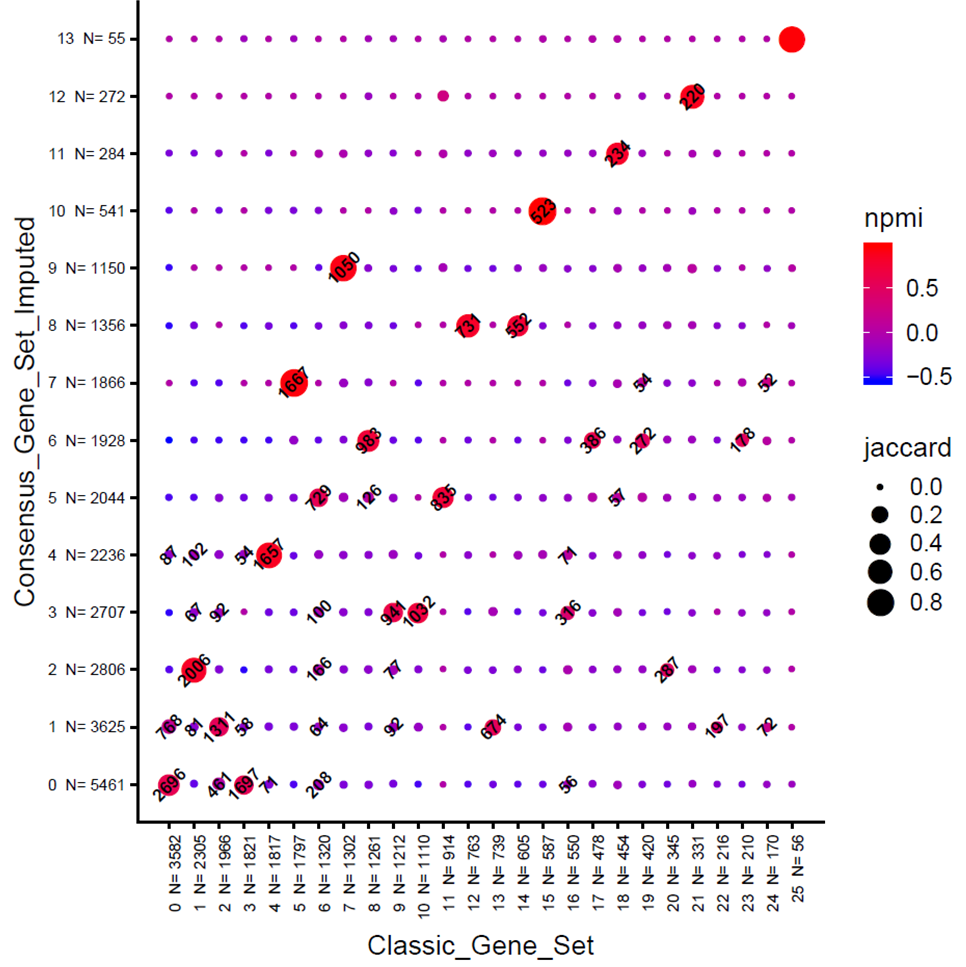

**Figure S6**

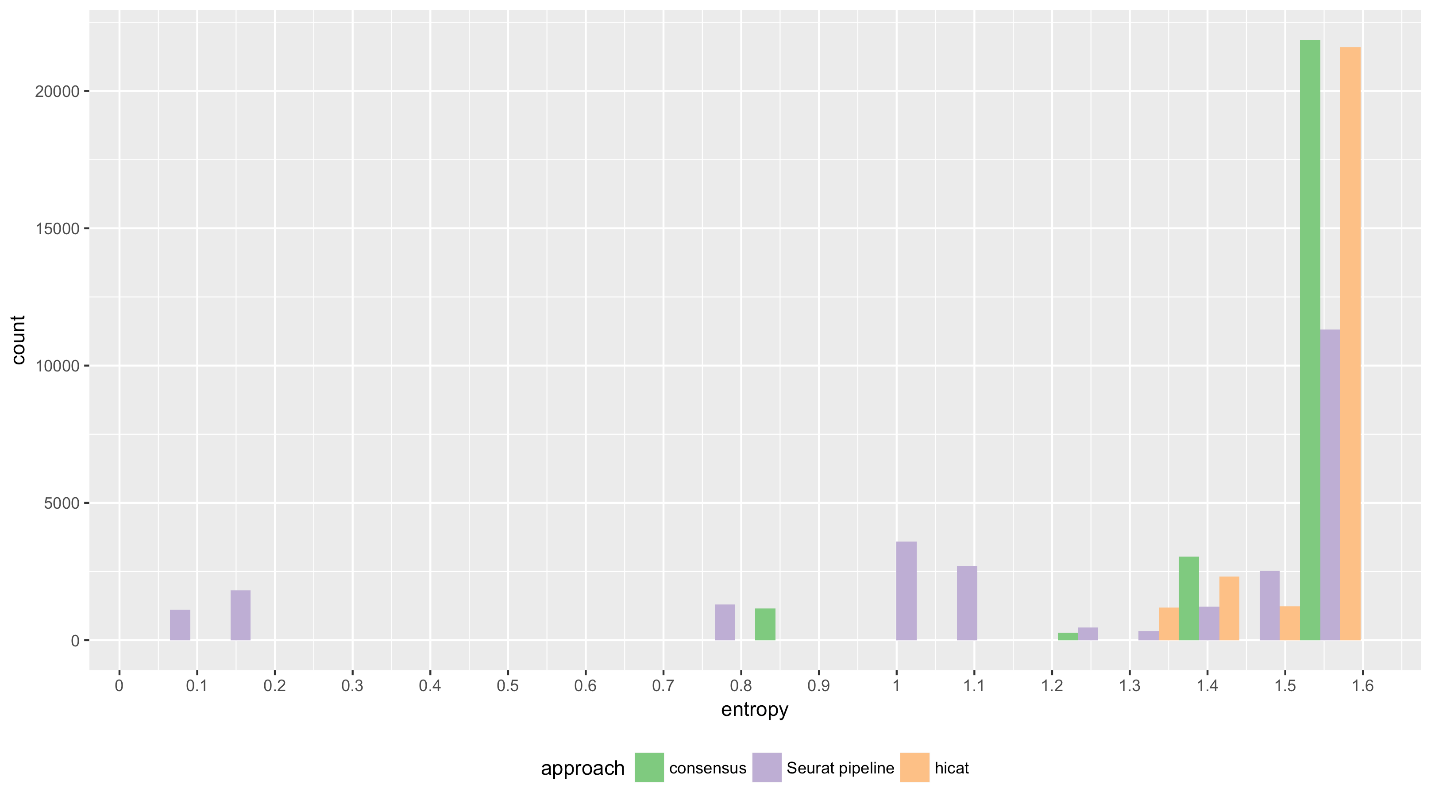

**Figure S7**

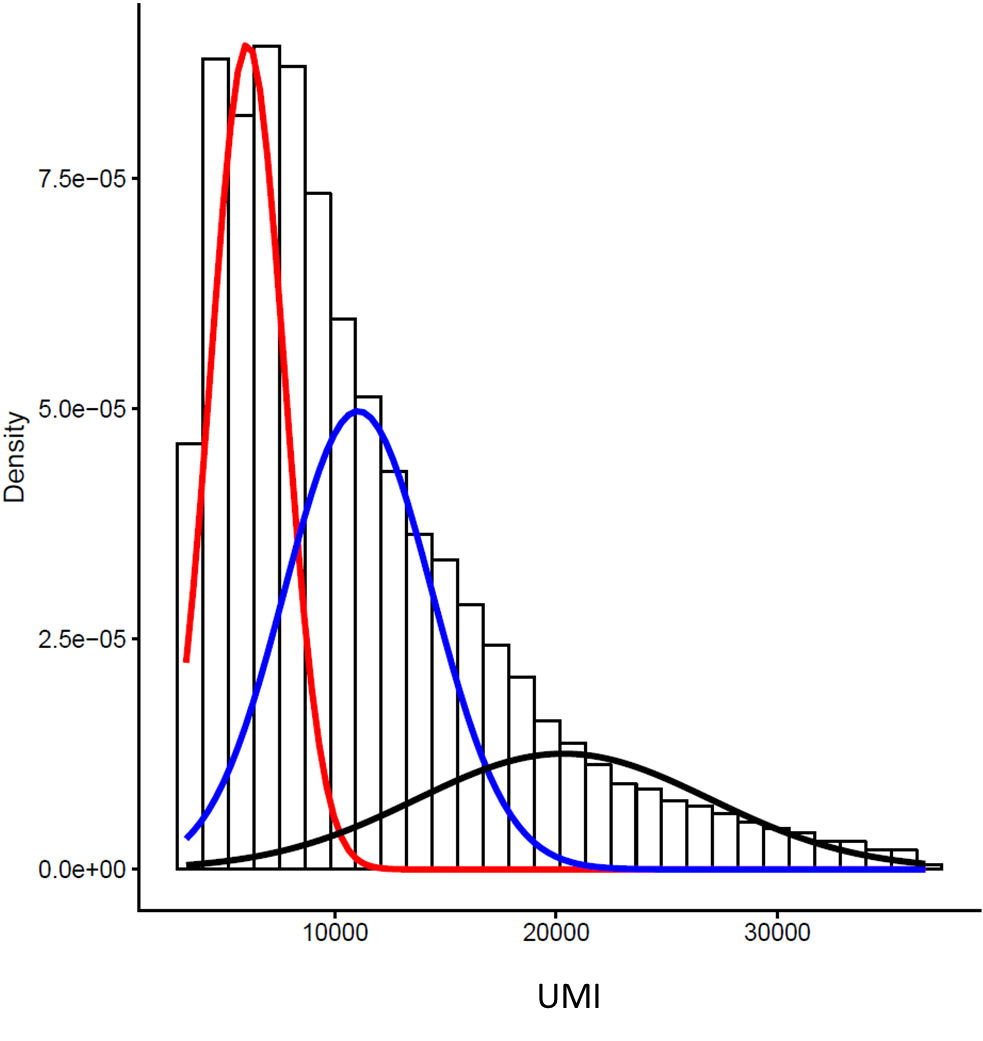

**Figure S8**
